# On-chip brain slice stimulation: precise control of electric fields and tissue orientation

**DOI:** 10.1101/2023.04.13.536696

**Authors:** Sebastian Shaner, Han Lu, Maximilian Lenz, Shreyash Garg, Andreas Vlachos, Maria Asplund

**Affiliations:** Department of Microsystems Engineering, University of Freiburg, Freiburg, Germany; BrainLinks-BrainTools Center, University of Freiburg, Freiburg, Germany; Department of Neuroanatomy, Institute of Anatomy and Cell Biology, Faculty of Medicine, University of Freiburg, Freiburg, Germany; MSc Neuroscience Program, Faculty of Biology, University of Freiburg, Freiburg, Germany; Center for Basics in Neuromodulation (NeuroModulBasics), Faculty of Medicine, University of Freiburg, Freiburg, Germany; Freiburg Institute for Advanced Studies (FRIAS), University of Freiburg, Freiburg, Germany; Division of Nursing and Medical Technology, Luleå University of Technology, Luleå, Sweden; Department of Microtechnology and Nanoscience, Chalmers University of Technology, Gothenburg, Sweden

**Keywords:** non-invasive brain stimulation, transcranial direct current stimulation, direct current electric field, bioelectronics, microfluidics, conducting hydrogels

## Abstract

Non-invasive brain stimulation modalities, including transcranial direct current stimulation (tDCS), are widely used in neuroscience and clinical practice to modulate brain function and treat neuropsychiatric diseases. DC stimulation of *ex vivo* brain tissue slices has been a method used to understand mechanisms imparted by tDCS. However, delivering spatiotemporally uniform direct current electric fields (dcEFs) that have precisely engineered magnitudes and are also exempt from toxic electrochemical by-products are both significant limitations in conventional experimental setups. As a consequence, bioelectronic dose-response interrelations, the role of EF orientation, and the biomechanisms of prolonged or repeated stimulation over several days all remain not well understood. Here we developed a platform with fluidic, electrochemical, and magnetically-induced spatial control. Fluidically, the chamber geometrically confines precise dcEF delivery to the enclosed brain slice and allows for tissue recovery in order to monitor post-stimulation effects. Electrochemically, conducting hydrogel electrodes mitigate stimulation-induced faradaic reactions typical of commonly-used metal electrodes. Magnetically, we applied ferromagnetic substrates beneath the tissue and used an external permanent magnet to enable *in situ* rotational control in relation to the dcEF. By combining the microfluidic chamber with live-cell calcium imaging and electrophysiological recordings, we showcased the potential to study the acute and lasting effects of dcEFs with the potential of providing multi-session stimulation. This on-chip bioelectronic platform presents a modernized yet simple solution to electrically stimulate explanted tissue by offering more environmental control to users, which unlocks new opportunities to conduct thorough brain stimulation mechanistic investigations.

**Graphical abstract:** 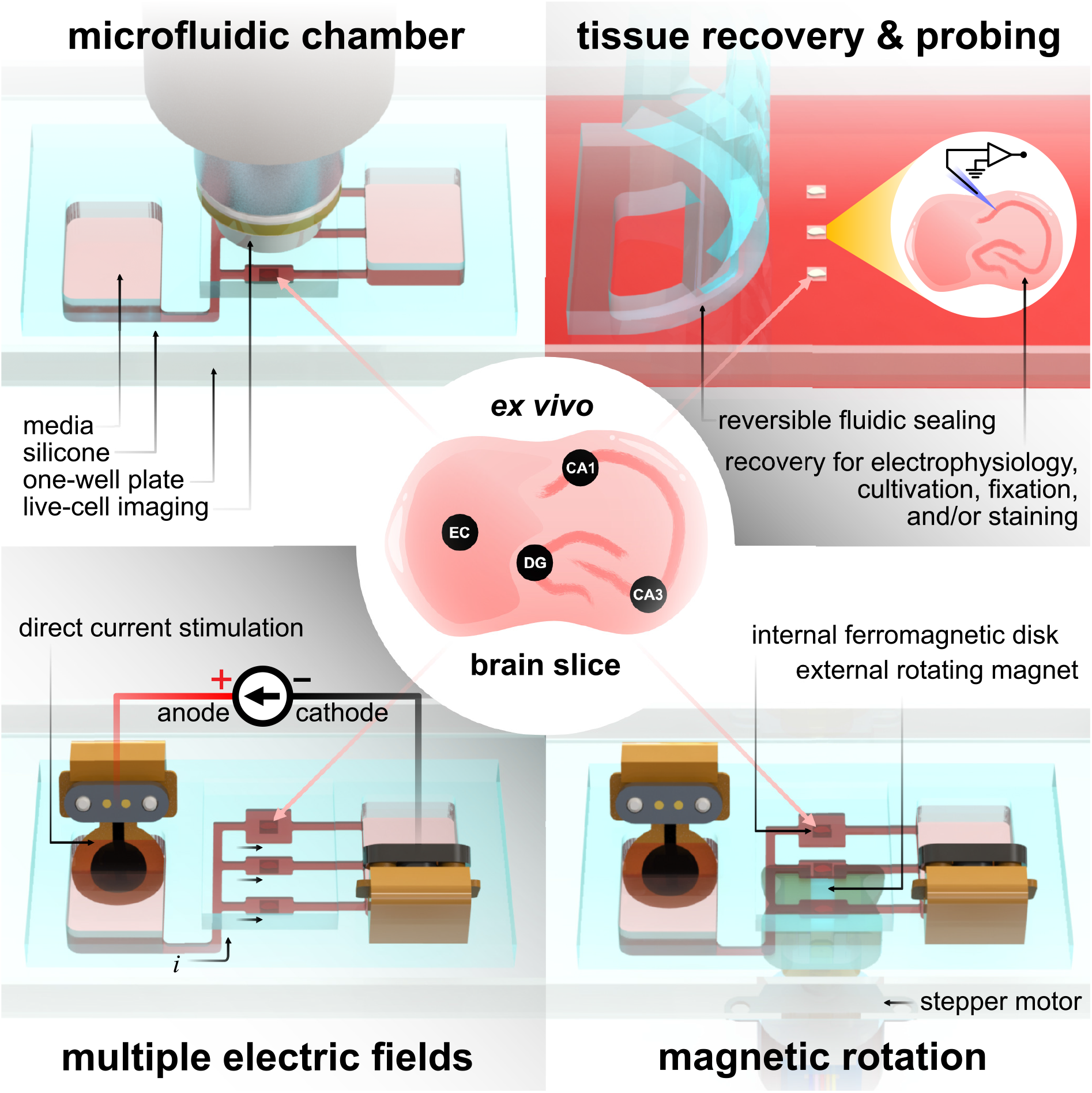

## Introduction

Microfluidic-based devices (*e.g.,* lab-on-a-chip) have been pivotal for advancing biomedical analysis from the protein to the organ level.^1, 2^ Neuroscientists have leveraged this technology to study numerous neural phenomena. This includes monitoring neural development and its behavioral relevance in small animal models *in vivo* (*C. elegans*, fruit flies, and zebrafish)^3, 4^ or investigating neural responses to external electrical, mechanical, or chemical stimuli^5, 6^ in *ex vivo* human and rodent brain tissue.^7, 8^ Additionally, *in vitro* cultured brain organoids or brain-on-a-chip models are frequently used to reduce or replace animal experimentation.^9–11^ Compared to traditional *in vivo* and *in vitro* approaches, the microfluidic regime possesses inherently reduced spatial dimensions that enable precise control of the microenvironment with low reagent consumption and high customizability via established microfabrication technologies.^12^ Specifically, microfluidic devices provide an alternative *in vitro* system with possibilities to better recapitulate the dynamic fluidic and biochemical environment in actual brains to study the neural mechanisms in large mammals where *in vivo* probing for broad parametric investigations are limited by ethical considerations, such as in human brains.^7, 13^ These on-chip platforms can facilitate fundamental discoveries for applications that require meticulous experimental tuning, such as brain stimulation.

Non-invasive brain stimulation is a promising approach for treating neuropsychiatric diseases,^14–16^ facilitating post-stroke rehabilitation,^17, 18^ and modulating learning and memory in humans.^19, 20^ Common stimulation modalities include repetitive transcranial magnetic stimulation (rTMS) and transcranial alternating or direct current stimulation (tACS/tDCS).^21^ Further optimizing their clinical stimulation protocols requires understanding the action mechanisms of induced or applied electric fields (EFs), especially the impact of direct current electric fields (dcEFs) on neural activity. This is not only pivotal for tDCS applications but also essential in juxtaposing the effects produced by alternating EFs that are seen in tACS and rTMS.

*In vitro* tDCS studies, which will be referred to as (t)DCS studies in this manuscript, on both the single neuron and neural network levels have shown weak EFs at subthreshold magnitudes can elevate neural membrane potential to shorten spike timing and increase the firing rate of an active neuron.^22–25^ The EF intensity,^23, 26^ the relative orientation between EF and the neural somato-dendritic axis,^27–30^ as well as the background activity level of the neural network,^31^ all lead to different extents of membrane potential polarization and the yielding aftereffects. Most of these *in vitro* studies were performed in acute brain slices on electrophysiological setups, where a fluidic perfusion system was integrated to maintain cell viability and parallel silver-silver chloride (Ag/AgCl) electrodes were adopted to generate the EF. Our recent review scrutinized these sophisticated setups and revealed their inability to precisely control the electric field (EF) and the risk of negative DC stimulation by-products.^32^ The short lifetime of acute slices prevents researchers from studying the effects of repetitive EF stimulation (*e.g.,* several sessions per day) or revisiting the neural tissue hours or days after EF stimulation. Some studies have attempted to address this by placing parallel electrodes inside six-well plates to stimulate tissue or cells inside the incubator,^33, 34^ while a lack of precise EF control and inevitable electrochemical faradaic reactions still remained. Other researchers adopted a widespread approach used in electrotaxis experiments where tissue or cells are surrounded with rectangular fluidic enclosures and are electrochemically connected to stimulation electrodes via agar or agarose salt bridges.^35^ However, the use of Ag/AgCl electrodes for direct current (DC) stimulation can lead to significant toxic Ag^+^ elution from the anode that travels nearly 1 cm per Coulomb of transferred charge.^36, 37^ In addition to the dueling merits and shortcomings of these platforms, being cumbersome and convoluted has limited the reproducibility of (t)DCS works. Therefore, a more customizable, affordable, and convenient alternative is needed.

We present here a microfluidic device for high throughput precision experiments with reversible fluidic sealing that can reliably investigate (t)DCS dose-response mechanisms. Our platform offers three notable advancements over standard (t)DCS platforms: (1) practical yet precise control of dcEF intensity by means of a microfluidic architecture, (2) reduced negative impact of electrochemical faradaic reactions via improved electrode materials, and (3) external control of tissue rotation relative to the dcEF while inside the chamber. This work first sets out to explain the design of the microfluidic chamber that facilitates efficient EF delivery while also minimizing faradaic reactions by using supercapacitive conducting hydrogel electrodes. A current divider microchannel network allows for the simultaneous stimulation of multiple brain slices at distinct EF intensities with a single input current. Additionally, a tissue-bound ferromagentic substrate allows for non-contact tissue orientation within the stimulation chamber, allowing investigation of tissue-EF orientation activity dependencies. Apart from the technical calibration, we also utilized organotypic entorhinal-hippocampus tissue cultures to confirm that the device does not affect cell viability, cause immune responses, or alter the cell membrane properties and neural synaptic transmission via a variety of staining, imaging, and electrophysiological techniques. Particularly, we showcased the possibility of using calcium imaging to assess the effects of dcEF at the network level during the stimulation. A visual overview of the platform technology and configuration options can be found in Fig. 1, which also serves as an experimental setup guide for all the figures in this manuscript.

**Fig. 1:**
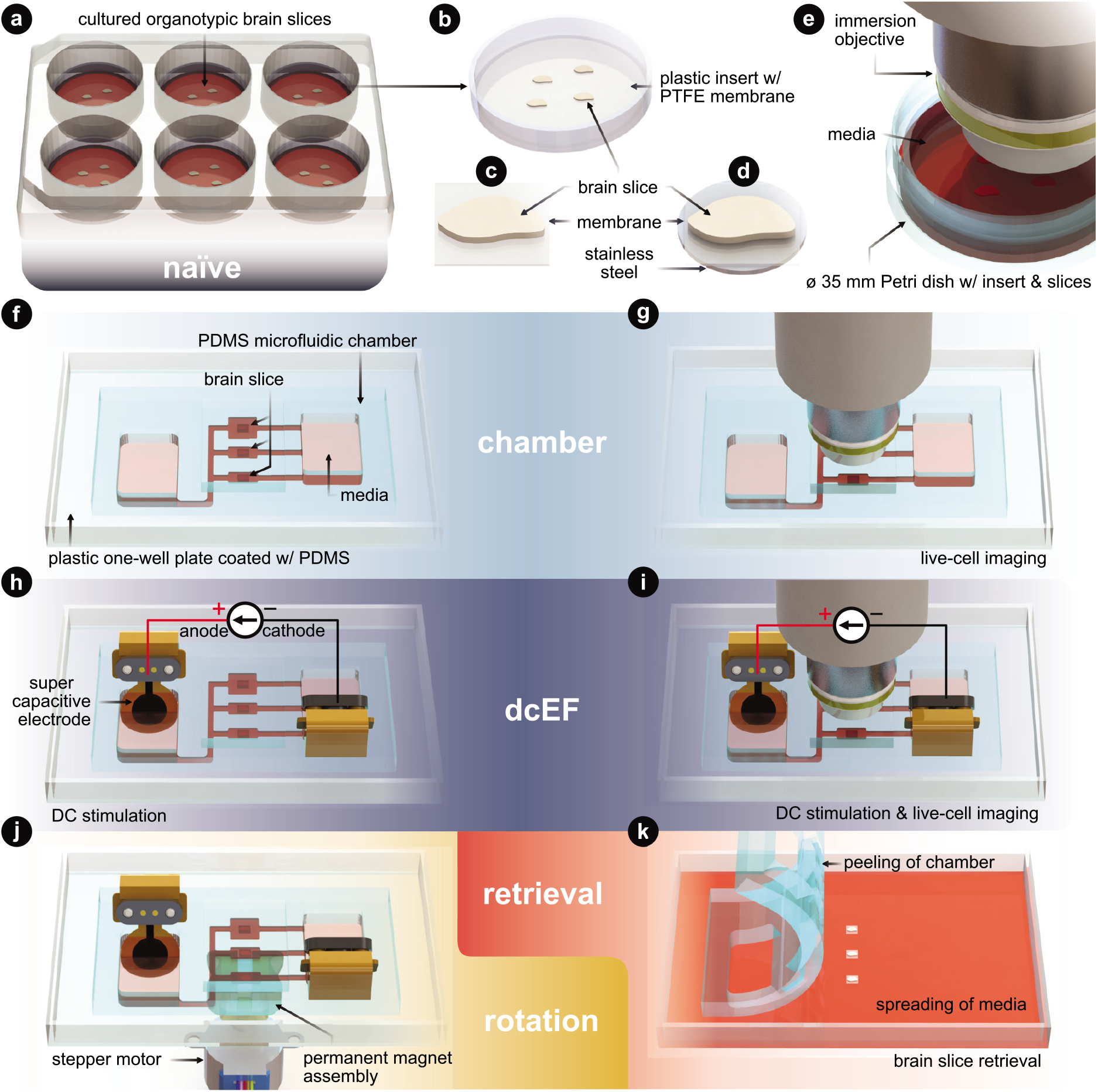
Technology and configuration features. (a) The *naïve* condition entails organotypic brain slices that are cultured on inserts. These inserts sit within a six-well plate and are maintained in an incubated environment. The slices are cultured with an air-liquid interface. They receive the media’s nutrients from below, while the top of the slices is exposed to the humidified air. (b) Plastic-framed insert with brain slices integrated onto a polytetrafluoroethylene (PTFE) membrane. (c) An individual brain slice that has been cut from the insert. (d) An individual brain slice that had stainless steel disk glued to the bottom of the membrane and then cut from the insert. (e) Live-cell imaging of the slices in a standard Petri dish. (f) The *chamber* condition where individual slices are assembled into the silicone (polydimethylsiloxane - PDMS) microfluidic chamber. (g) Live-cell imaging of slices within the microfluidic chamber. (h) The *dcEF* condition where individual slices are assembled into the microfluidic chamber and stimulated with supercapacitive electrodes to generate precise direct current electric fields (dcEFs). (i) Live-cell imaging and dcEF of slices within the microfluidic chamber. (j) Rotational control of the slice using (d) and a programmable rotating permanent magnet assembly. (k) Chamber is peeled and slices are be retrieved for further *post hoc* assessments. Fig. 3 uses (a),(f),(h). Fig. 4 uses (e),(g),(i),(k). Fig. 5 uses (a),(f),(h),(k). Fig. 6 uses (e),(g),(i). Fig. 7 uses (h),(j),(k).

## Results

### Microfluidic chamber generates spatiotemporally uniform direct current EFs

DC stimulation of explanted brain tissue is conventionally carried out in an open bath within electrophysiological setups.^23, 29, 38^ However, generating spatiotemporally uniform dcEFs with precise EF intensity control in these setups is difficult (Supplemental Fig. S1).^32^ Meanwhile, this control is fundamental to studying dose-dependent and EF orientation-related neural mechanisms. Therefore, we leveraged the microchannel design used in cell electrotaxis experiments^39, 40^ to create a microfluidic chamber with geometric confinement around the brain tissue, where the controlled volume allows for precise delivery of dcEFs in and around the tissue (Fig. 2). The advantage of the four-walled rectangular confinement around the tissue is that the dcEF within the microchannel can be easily calculated through the application of Ohm’s and Pouillet’s laws (*E* = *i/*(*σwh*) in V m*^−^*^1^) with the known material properties of the electrolyte (electrical conductivity, *σ* in S m*^−^*^1^), geometric properties of the channel (cross-sectional area, *A* = *w · h* in m^2^), and the constant input stimulation current (*i* in A).

**Fig. 2:**
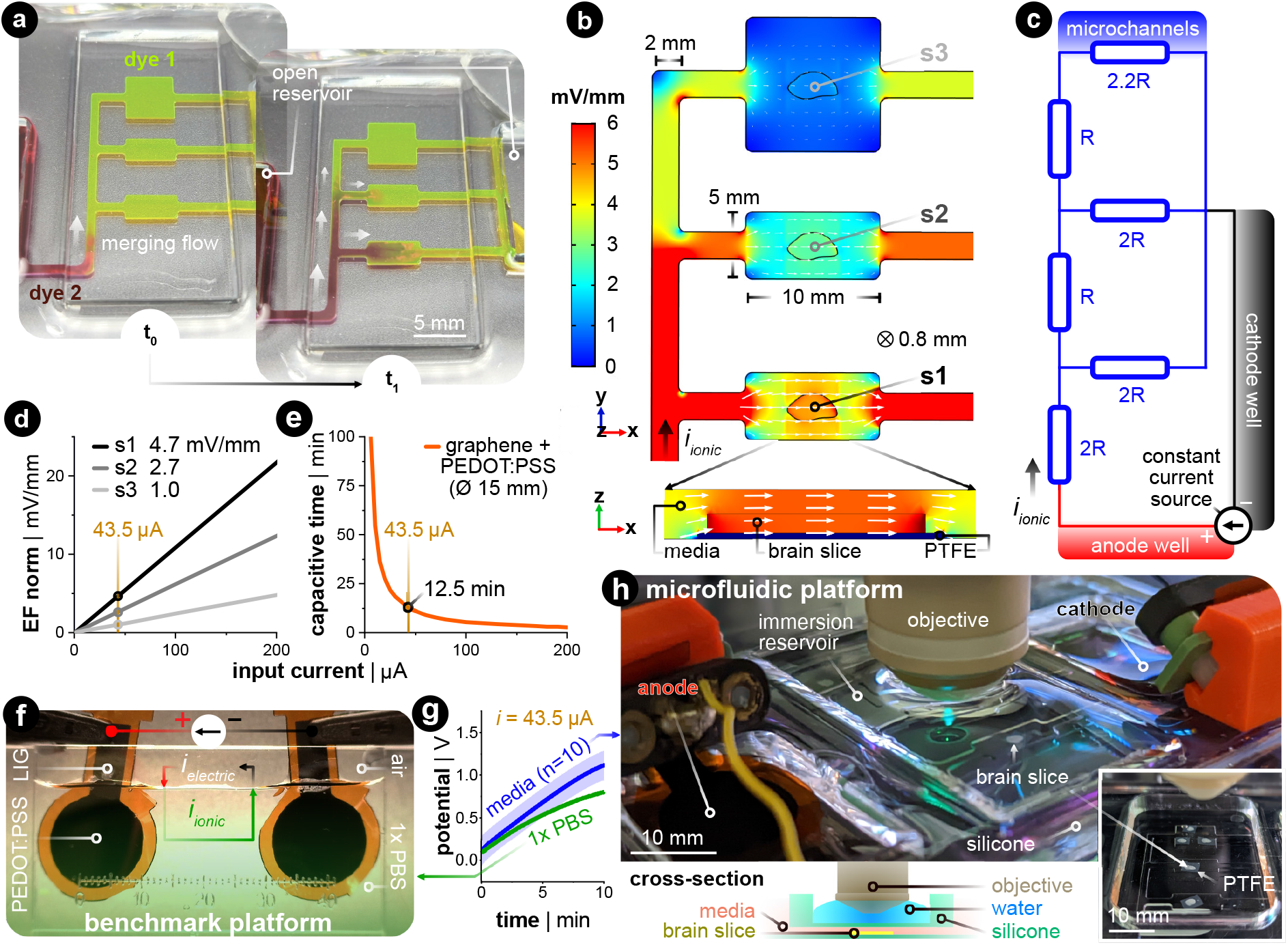
Controlling the electric field dosage in a microfluidic current divider chamber. (a) Primed microfluidic chamber with dye 1 (yellow) was spiked with dye 2 (red) to visually analogize fluidic resistivity to electrical resistivity. *t*_0_ shows when dye 2 was added to the open reservoir on the left. *t*_1_ shows a few seconds later where the merging flow branches into the current divider network. (b) FEA simulation of the EF magnitude and distribution in the parallel microchannels with a given constant input current (*i*_ionic_ = 43.5 µA). Note that the *x*- and *y*-dimensions of the design are annotated in the figure, where the thickness (*z*-dimension) is 800 µm. s1, s2, and s3 are three brain slices submerged in the microchannel. The white arrow’s orientation and size signify EF direction and intensity in the stimulation zone, respectively. The *x*,*y*-plane is at the tissue’s mid-plane (*z* = 400 µm). The magnified cross-sectional view shows s1’s *xz*-plane. (c) The equivalent electrical circuit of the current’s path throughout the electrolyte-filled microfluidic network. The parallel branches of ‘2*R*” and ‘2.2*R*” constitutes the summation of the smaller and wider channels in the *x*-direction. (d) Relationship between the input current and the generated EF within brain slices in the three parallel stimulation zones. (e) Estimation of capacitive discharge current time based on the input current when using large conducting hydrogel (PEDOT:PSS-coated LIG) electrodes. The yellow reference lines in (d) and (e) indicate the current used throughout the rest of this work. (f) Constant current benchmarking setup of the PEDOT:PSS-coated LIG electrodes in 1*x* PBS. The green shaded area shows the ionic current regime. (g) Voltage excursion data when using PEDOT:PSS-coated LIG electrodes in two different electrochemical setups, both with a constant current of 43.5 µA. The test setup (in green and panel f) does not have a microchannel, while the final setup (in blue and panel h) is the microfluidic assembly. Both cases show a capacitive-dominant charging profile. (h) Final configuration of the stimulation chamber. The complete silicone chamber has a built-in reservoir above the microchannels for an immersion-based objective used in live-cell confocal imaging. The cross-section illustration shows how the silicone separates the media from the immersion imaging reservoir.

In order to allow for concurrent stimulation of multiple brain slices, we specifically leveraged a current splitting design with three parallel microchannels where each channel could accommodate one brain slice. Each slice is subjected to a distinct EF strength, which all stem from a single pair of electrodes. As the current travels from the anode to the cathode, it splits at each bifurcating junction such that the EF is lowered as the current traverses through the device. As an illustration, dyes were used to visualize the microchannel network’s fluidic resistance as an analog to the electrical resistance^41^ (Fig. 2a). Specifically, the yellow-colored dye (fluorescein) was used to displace the air-filled channels and to fluidically prime the device, then the maroon-colored dye (Congo red) was added to the open inlet reservoir. Hydrostatic pressure-induced flow mixes the two dyes within the channels and indirectly illustrates the ionic current’s path throughout the microfluidic network, albeit at a much slower time scale (Supplemental Video 1). Given the precedent that finite element analysis (FEA) can predict the experimentally measured dcEFs in microchannels with considerable precision,^39^ FEA simulation was utilized to elucidate both the EF distribution and magnitude in our design for a given input current. Simulation results confirmed that the relative EF magnitudes resemble the merging flow dynamics (Fig. 2b). The submerged slice was also stimulated homogeneously throughout the tissue, which can be seen in the cross-sectional view of device section s1. An equivalent electrical circuit of the microfluidic network further encompasses how the current is divided in each parallel branch that houses the brain slices (Fig. 2c). Since the microfluidic geometry ensures control of the cross-section, the EF strength scales linearly with the input current (Fig. 2d).

The stimulation electrode material is another factor to consider for delivering constant DC with minimal electrochemically-generated species, which are side reactions to the DC.^32, 42^ Here, non-metal supercapacitive electrodes (laser-induced graphene coated with the conducting hydrogel PEDOT:PSS) were used.^43^ Slow cyclic voltammetry (CV) was used to assess the capacitance of the electrode (Supplemental Note S1 and Fig. S2). This capacitance value was then used to estimate the electrode’s capacitive discharge current as a function of a given constant input current (Fig. 2e). These values were then used to determine the electrode size needed for the EF magnitudes included in this study. The rationale for the EF intensity range used in this study is to replicate values reported to be efficient in previous human (*<* 1 mV mm*^−^*^1^)^44^ and *in vitro* studies (*<* 5 mV mm*^−^*^1^).^16, 45, 46^ To have all three stimulation zones within this range, we chose an input current value of 43.5 µA for the 15 mm diameter hydrogel electrodes.

This input current generates three representative EF intensities, 4.7, 2.7, and 1.0 mV mm*^−^*^1^ in sections s1, s2, and s3, respectively (Fig. 2d). This allows for up to 12.5 min of capacitive-dominant current (Fig. 2e). We chose 10 min as the standard stimulation duration, which is also within the clinical stimulation time frame.^47^

Finally, we compared the PEDOT:PSS hydrogel-coated LIG to common electrode materials used in (t)DCS studies: Ag/AgCl and platinum (Pt). We applied constant monophasic DC for the same duration and current density for all three electrode materials in a two-electrode setup with a physiologically-relevant electrolyte (Supplemental Fig. S3). This benchmarking setup electrochemically mimics the final microfluidic chamber, as the 1*x* phosphate-buffered saline (PBS) has the same ionic conductivity (1.5 S m*^−^*^1^) as the incubation media used throughout the rest of this work (Fig. 2f). The monitored voltage excursion demonstrates that the non-polarizable Ag/AgCl electrode predominately transferred charge via faradaic reactions. The Pt electrode quickly discharged the capacitive electrochemical double layer (ECDL) and subsequently transitioned into a hydrogen adsorption-induced pseudocapacitive current^48, 49^ and faradaic current. In contrast to both metal electrodes, the PEDOT:PSS hydrogel electrode exhibits a capacitive discharge current due to its large volumetric storage of ionic charge (Fig. 2g in green, Supplemental Fig. S3 in orange). These electrodes were also tested with the same input current in the final microfluidic platform, but now with culture media as the electrolyte (Fig. 2g in blue). The potential rises quicker for the final platform due the introduction of the microfluidic resistor network. However, the linear slope for both electrochemical setups suggests that current is predominately, if not entirely, provided in a capacitive manner. By staying within the time of capacitive discharge, pH shifts and generation of reactive oxygen species (ROS) via faradaic reactions are kept to a minimum.^50^ The complete fluidic and electrical package fits within a standard one-well plate and can be used with either upright or inverted microscopes (Fig. 2h, Supplemental Fig. S4).

### Chamber exposure did not harm cell viability nor cause immune responses in tissue culture

In order to probe whether the silicone microfluidic chamber would affect cell viability, tissue cultures prepared from wild-type mice were placed into the sealed chamber for 20 min and stained for dead cell nuclei with SYTOX-green (Fig. 3a). As summarized in Fig. 3b-c, immersion inside the microfluidic chamber for 20 min did not significantly increase the SYTOX-green signal intensity in comparison to näıve cultures (Kruskal-Wallis test, *p >* 0.99). Thus, this attests that our fluidic system is compatible with brain slice tissue cultures. Visual inspection of DAPI stained cells furthermore confirmed this result (in small insets and in Supplemental Fig. S5). We furthermore included positive control cultures that were treated with 50 *µ*M NMDA for four hours, and this treated group showed significantly higher SYTOX-green signal intensity compared to näıve cultures (Kruskal-Wallis test, *p* = 0.03), speaking to the prevalent excitotoxicity-induced cell death. Visual inspection of the DAPI signal also confirmed condensed nuclei shape in CA1 of NMDA-treated cultures, which was absent in chamber-exposed and näıve cultures. The SYTOX-green signal intensity of the cultures being stimulated by weak dcEF (4.7 mV mm*^−^*^1^) showed a reduction tendency but was not significantly different from näıve cultures or chamber-treated cultures (Kruskal-Wallis test, *p* = 0.06); DAPI signal was also normal in this group (Supplemental Fig. S5). Based on this, we conclude that immersion inside the silicone chamber for around 20 min with or without dcEF does not harm cell viability and that the presented platform is tissue compatible.

**Fig. 3:**
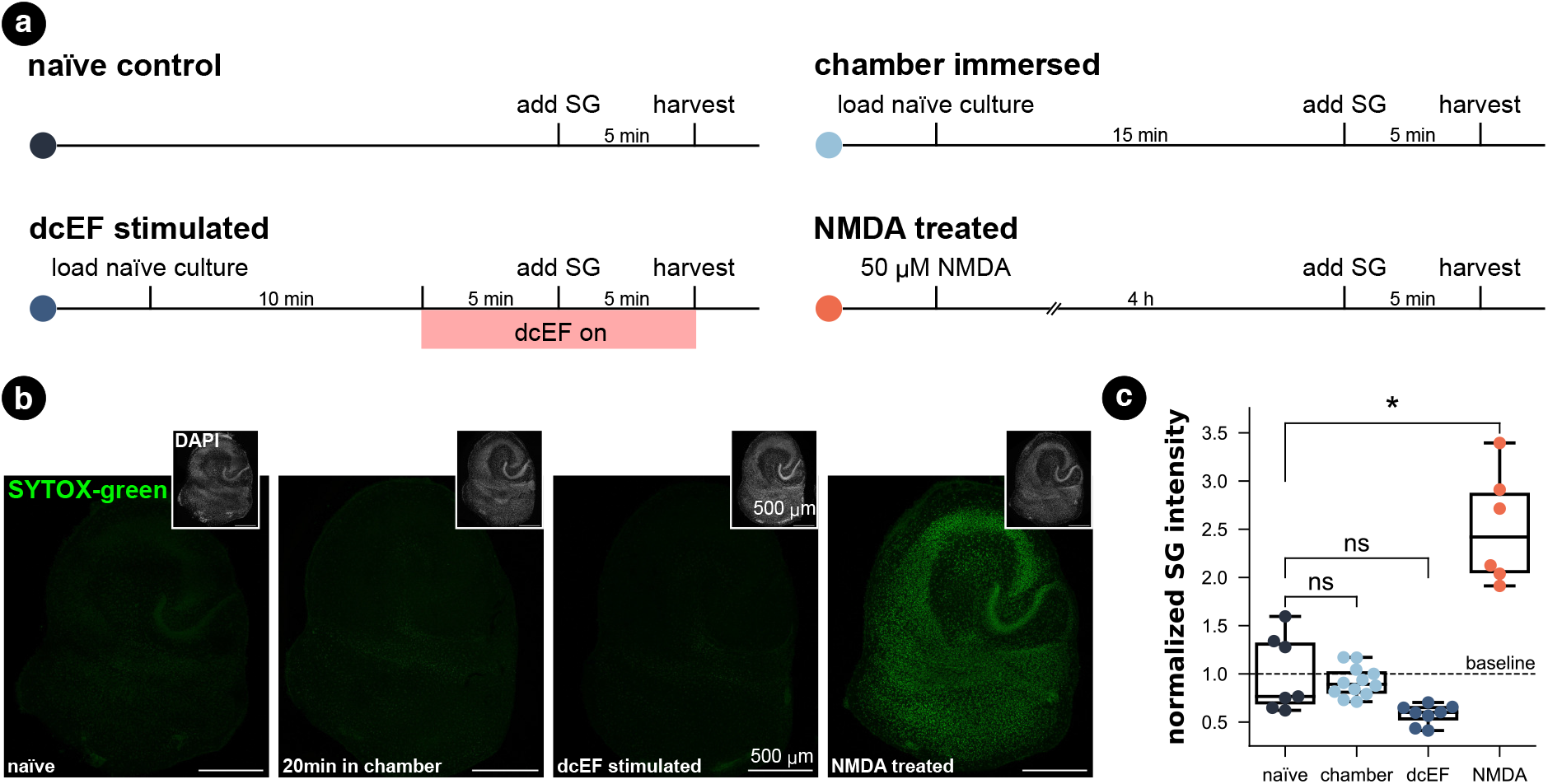
Tissue culture cell viability was maintained in both the microfluidic environment and after weak dcEF. (a) Experimental design for four groups: the näıve control, the chamber immersion group where cultures were immersed inside the chamber for 20 min without dcEF, the dcEF stimulation group with 4.7 mV*/*mm, the positive control where cultures were treated with 50 *µ*M NMDA for four hours. (b) Representative images of SYTOX-green (SG) staining in four groups. The small insets display the DAPI signal of the corresponding cultures. (c) Normalized SYTOX-green (SG) signal intensity of the whole culture in four groups (*N* = 7 for the näıve control; *N* = 12 for chamber immersion group; *N* = 8 for dcEF stimulation group; *N* = 6 for NMDA-treated positive control group). The raw values were normalized by the mean value of the näıve control group. Box plots summarize the mean, quartiles, and distribution of each condition. The Kruskal-Wallis test followed by Dunn’s multiple comparisons test was used for group and pairwise comparisons. If not otherwise stated, *∗* is *p <* 0.05, *∗∗* is *p <* 0.01, *∗ ∗ ∗* is *p <* 0.001, while “ns” means *p >* 0.05. Scale bars equal to 500 *µ*m.

In addition, we investigated for inflammation by using live-cell imaging to examine tissue cultures prepared from a transgenic mouse line, which expresses enhanced green fluorescent protein (eGPF) under the promoter of the inflammatory cytokine tumor necrosis factor alpha (TNF*α*).^51^ The eGFP signal intensity that reflects TNF*α* expression was imaged before and after each treatment (Fig. 4a-c). Consistent with our SYTOX-green results, neither chamber exposure nor dcEF stimulation (4.7 mV mm*^−^*^1^) increased the eGFP signal intensity (Wilcoxon test, *p* = 0.10 and *p* = 0.29, respectively). As previously shown,^51^ three-day bacterial lipopolysaccharide (LPS) (1 *µ*g*/*mL) treatment, which is known to trigger inflammation, induced a significant increase in eGFP signal (*i.e.,* expression of TNF*α*) in the positive control cultures (Wilcoxon test, *p* = 0.0004). In summary, we conclude that immersion or stimulation of cultures inside the silicone chamber does not harm cell viability nor cause overt immune responses. Thus, the methodological benefits of the microfluidic environment come at no cost in terms of tissue viability.

**Fig. 4:**
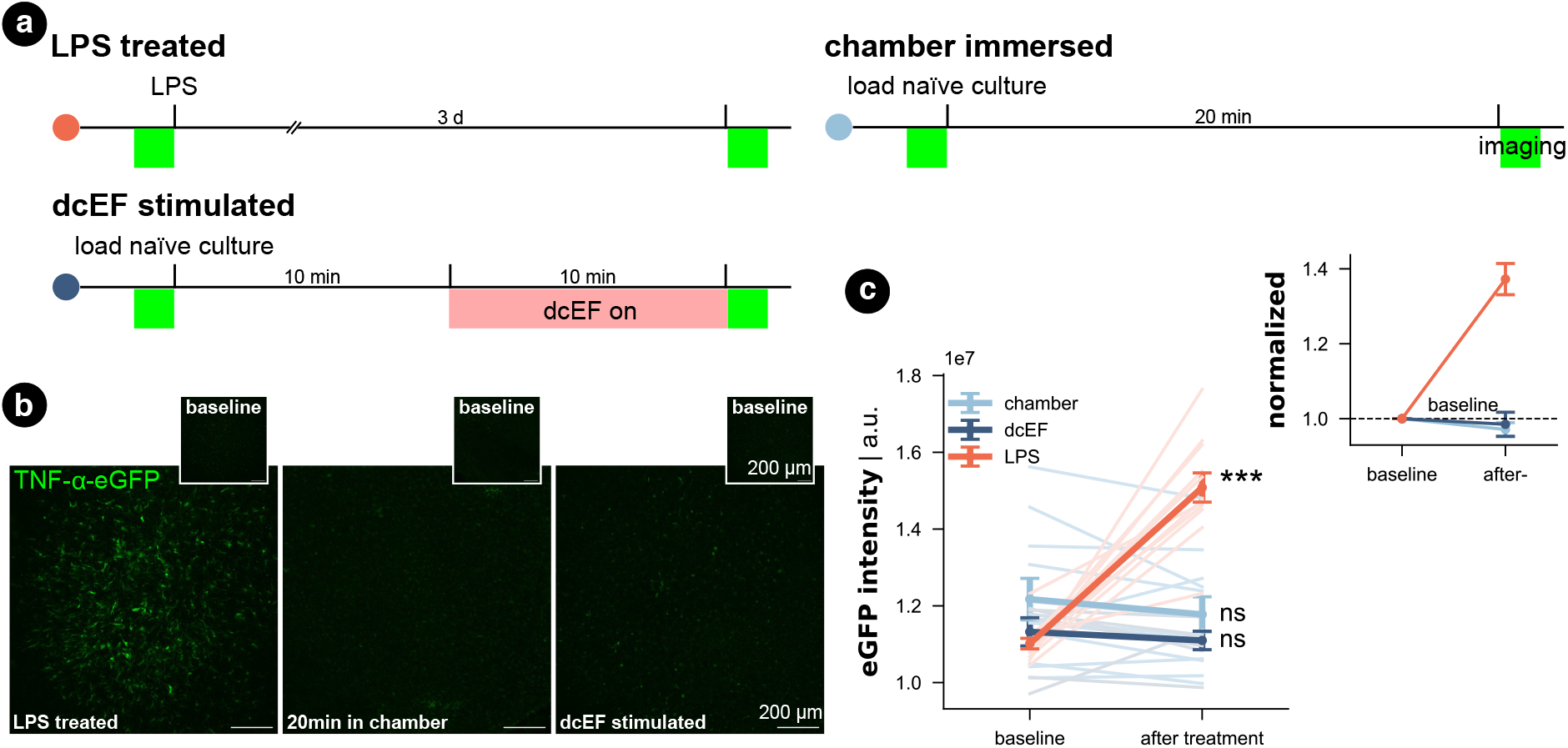
Inflammatory response did not occur for tissue cultures in either the microfluidic environment or after weak dcEF. (a) Experiment design for three groups. (b) Representative images of TNF*α*-eGFP culture before and after corresponding treatment: three-day LPS treatment (1 *µ*g*/*mL); 20 min immersion inside the chamber with or without 10 min dcEF stimulation at 4.7 mV*/*mm. (c) Quantified eGFP signal intensity of the whole culture for the three experimental groups. An arbitrary unit (a.u.) was used. The lines with light shades are raw data for individual cultures while the three lines with strong shades are averaged data for each group with standard error of the mean (s.e.m.) as the error bar (*N* = 11 for the chamber immersion group; *N* = 7 for the dcEF stimulation group; *N* = 12 for the LPS-treated group.) The inset displays the averaged values normalized by the corresponding baseline intensity of individual cultures. Wilcoxon test was used for statistical analysis of each group. Scale bars equal to 200 *µ*m.

### Synaptic transmission and intrinsic membrane properties remained intact in the chamber-housed neurons

Whole-cell patch-clamp recordings were performed on cultures immediately after 20 min of immersion to assess whether the handling and chamber immersion altered the neural functional properties compared to näıve cultures. Since whole-cell patch-clamp recording is widely used to probe synaptic plasticity, we also have the chance to examine the aftereffects of dcEF stimulation with the same measurements. Therefore, we stimulated cultures as previously described (4.7 mV mm*^−^*^1^) and re-cultured them back in the incubator for around 2 *±* 1 h before recording. This is to allow enough time for synaptic plasticity induction.

A separate group of chamber-immersed cultures that underwent the same retrieving and re-culturing procedure served as chamber controls (Fig. 5a). We examined the excitatory synaptic transmission and the intrinsic membrane properties of CA1 pyramidal neurons in these experiments (see the representative image in Fig. 5b). In line with our SYTOX-green results and TNF*α*/eGFP results, visual inspection did not indicate cell death in any of the groups. Also, the whole-cell configuration of all recorded neurons was equally well established. No significant differences in the mean amplitude, half-width, and frequency of AMPA-receptor-mediated spontaneous excitatory postsynaptic currents (sEPSCS) were observed among the four groups (Kruskal-Wallis test, *p >* 0.05 for group level and pair-wise examinations, Fig. 5b-e). This suggests neither chamber immersion nor delivering stimulation inside the chamber altered neural synaptic transmission. Similarly, input-output curve analysis showed no significant difference in resting membrane potential among the four groups (Kruskal-Wallis test, *p >* 0.05 for group level and pair-wise examinations, Fig. 5f, left panel). However, a significant reduction in input resistance was observed in the two groups that underwent the 2 h retrieving and re-culturing procedure (Mann Whitney U test after grouping the data, *p <* 0.001, Fig. 5f, right panel). The action potential frequency analysis also shows that the act of re-culturing classified the four conditions into two categories. The chamber-immersed cultures showed similar kinetics in generating action potentials as näıve cultures. In contrast, both dcEF-stimulated and its chamber-control groups presented altered kinetics and required a larger magnitude of injection current for neurons to generate action potentials. Their elevated spiking frequency still persisted at higher current intensities; whereas, the other two groups of neurons displayed an attenuated tendency (see example traces in the four groups at 500 pA injection). Therefore, we concluded that mounting, immersing, and removing cultures from the silicone microfluidic chambers have no major effects on synaptic transmission or intrinsic membrane properties. Tissue cultures can be subjected to additional experimental assessment after stimulation, including methods sensitive to cell viability alterations, such as electrophysiological recordings. However, caution should be taken as re-culturing may amend neural response to external stimulation to a different level, such that one should always consider matching the corresponding control group. These control groups were used throughout the rest of this manuscript when re-culturing was needed for monitoring offline effects.

**Fig. 5:**
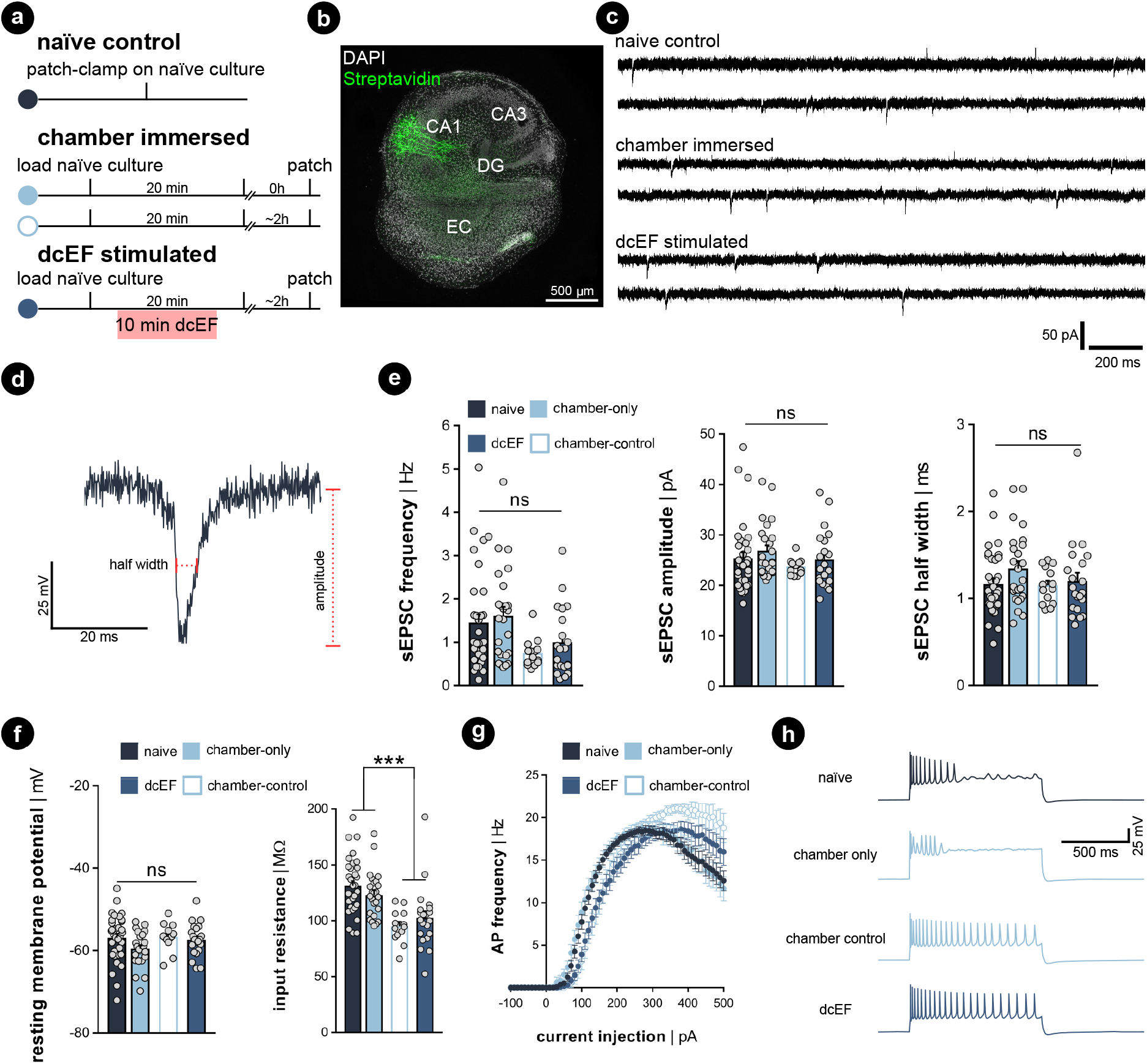
Excitatory synaptic transmission and passive membrane properties were not altered by the microfluidic environment. (a) Experimental protocol. (b) Example of recorded biocytin-filled CA1 pyramidal neurons. (c) Representative sEPSC traces recorded from näıve cultures, chamber- immersed cultures, and dcEF-stimulated cultures. (d) Amplified sample event and the measurement of sEPSC amplitude and half width. (e) Mean and s.e.m. of frequency, the average amplitude, and half width of sEPSCs recorded from individual neurons in four groups. Each dot represents one recorded neuron. *N* = 34 for näıve cells, *N* = 25 for chamber-only cells, *N* = 21 for dcEF cells, *N* = 14 for chamber control cells were recorded for all the analyses used in (e)-(g). (f) Mean and s.e.m. of resting membrane potential and input resistance of individual neurons in four groups. Each dot represents a neuron. Kruskal-Wallis test, followed by Dunn’s multiple comparisons test, was used for statistical analysis in (e) and (f). (g) Input-output curves of action potential frequency when different input current intensities were injected into the neuron. Each dot represents the averaged frequency, while the error bar indicates the s.e.m. across all recorded neurons. RM two-way ANOVA followed by Sidak’s multiple comparisons test was used for statistical analysis. No significant difference was detected between the chamber-only and näıve groups at all levels (*p >* 0.05); no significant difference was detected between the chamber-control and dcEF groups at all levels (*p >* 0.05). After merging the corresponding datasets, significant changes were detected at several levels between the groups with reculturing and without reculturing (*p <* 0.001). Scale bar equals to 500 *µm*.

### Calcium activity was preserved in chamber-immersed cultures and pulsed dcEFs increased calcium spikes

Since the whole-cell patch-clamp recordings did not show significant plasticity triggering effects with a weak dcEF at 4.7 mV mm*^−^*^1^, we performed calcium imaging to assess whether network activity was perturbed inside the chamber with and without stimulation. Therefore, we transfected wild-type cultures with AAV1-hSyn1-GCaMP6f-P2A-nls-dTomato virus (RRID:Addgene 51085) and simultaneously imaged multiple neurons in the CA1 region (Fig. 6a). We observed that when imaged inside the incubation medium, näıve cultures presented active calcium dynamics that the whole network synchronized at a low frequency where almost all neurons were activated. During the inactive status between synchronized network events, individual neurons varied in activity levels; some remained silent, some remained lit, while some fired sparse somata calcium spikes. We applied a computer vision algorithm and extracted the time series of multiple neurons per culture. A comparison of individual calcium traces obtained from the same culture showed both synchronized network events and neuron-specific events (red and yellow arrows in Fig. 6b). Despite the heterogeneity of calcium activity among neurons and cultures (Supplemental Fig. S6), our results demonstrated that immersion inside the chamber for 10 min reduced the overall calcium activity level (*p <* 0.001, unpaired student’s t-test; 99.99%CI = [9.09, 25.8], LLM), which agrees with our action potential frequency analysis. Further application of weak dcEF (4.7 mV mm*^−^*^1^) after a 10 min interval showed a temporal recovery of calcium activity, which was not attributed to or further boosted by dcEF stimulation (Supplemental Fig. S7a,b). In another set of cultures, we confirmed that weak dcEF stimulation at 4.7 mV mm*^−^*^1^ did not increase the expression of immediate early gene c-Fos either, which is a marker for neural activation (Supplemental Fig. S7c-d).

**Fig. 6:**
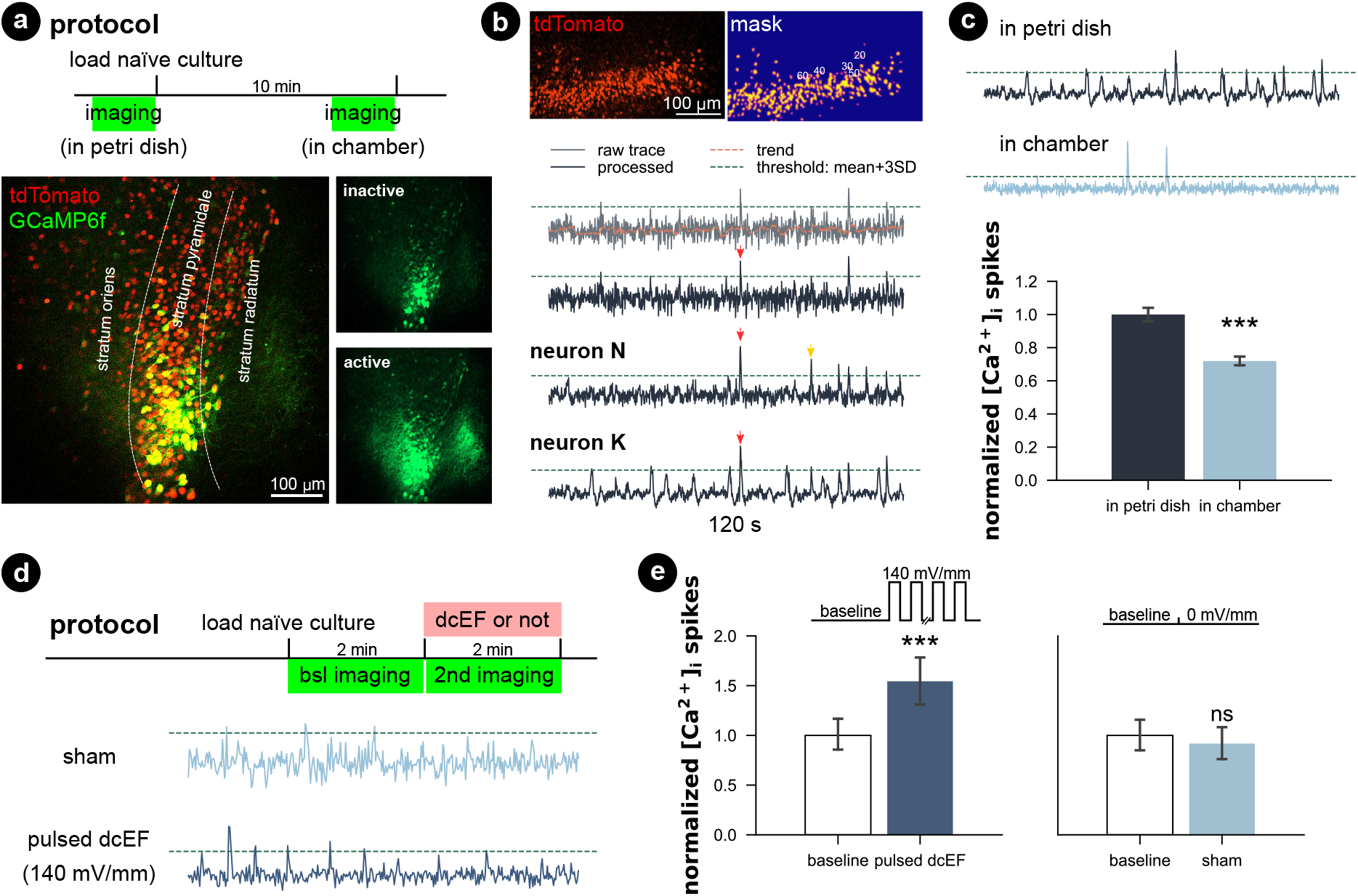
Calcium activity was preserved in microfluidic environment, while pulsed dcEF activated neurons. (a) The entire culture was transfected with AAV1-hSyn1-GCaMP6f-P2A-nls-dTomato virus. A within-subject design was applied, so each culture was imaged twice: inside a Petri dish before loading and after being immersed inside the chamber for 10 min. (b) Methods used to extract the time series of calcium signals of individual neurons and detect calcium spikes. Processed traces of three neurons obtained from the same culture showing both synchronized network events (red arrows) and neuron- specific events (yellow arrows). (c) Chamber immersion reduced the calcium activity. *N* = 14 cultures were used for imaging and each culture was imaged at both timings. *N* = 1153 neurons were identified in the Petri dish imaging phase, and *N* = 1002 neurons were identified in the chamber imaging phase. If not otherwise stated, the calcium spike rates were normalized by their corresponding average rate at baseline. (d) Protocol and example trace when we pulsed high dcEF to the immersed cultures. *N* = 4 cultures were used for the sham group (186 neurons identified at baseline and 178 neurons identified for during sham treatment), and *N* = 4 cultures were used for pulsed dcEF stimulation (235 neurons identified at baseline and 187 neurons identified for during stimulation). Each culture was imaged twice at baseline and under experimentation. (e) Strong pulsed dcEF stimulation immediately increased calcium spikes. Unpaired student’s t-test and linear mixed models grouped per culture were used for statistical analysis in (c) and (e).

To provide a clear effect of DC stimulation, a much higher dcEF (140 mV mm*^−^*^1^) was pulsed (100 on/off cycles: 0.1 s at 1.29 mA and 0.5 s interval) to the culture through agarose-embedded double-layered PEDOT:PSS electrodes (Supplemental Fig. S8), and an immediate increase in calcium spikes was observed (*p <* 0.001, unpaired student’s t-test; 99.99%CI = [9.09, 25.8], LLM; Fig. 6d-e). Note that this suprathreshold EF strength is close to those that are typically used in rTMS studies.^52, 53^ The specialized agarose-embedded double-layered PEDOT:PSS electrodes meant to further eliminate any potential electrochemical by-products from reaching the media (Supplemental Fig. S8).

Our results showcased that the platform preserves the calcium activity of submerged cultures. It is capable of monitoring the whole culture’s calcium activity and detecting both directional changes (increase or reduction) via non-contact imaging approaches throughout the experimentation. Similar to the electrophysiological experiments, we showed that handling and timing might set the calcium activity baseline differently, so as discussed in the previous section, a matched control group should be always carefully planned.

### Controlling brain slice orientation within the chamber via a non-contact magnetic approach

When modeling (t)DCS in cultures, it would be ideal if not only the intensity but also the orientation of the field could be precisely controlled.^54^ In conventional experimental settings, the orientation was studied by *post hoc* analysis of the morphology of different neurons^29^ by placing several groups of brain slices in correspondingly different orientations relative to the EF,^23^ or by studying different pathways.^55^ In these studies, one has to orientate the brain slice in the desired configuration before experiments, such as during the chamber assembly process. To enable changing culture orientation during experiments, we utilized a biocompatible, corrosion-resistant, and ferromagnetic grade of stainless steel (*i.e.,* 1.4310) to act as a rotational stage. The brain slice rests on the stage so that its orientation within the chamber can be controlled using an external permanent magnet (Fig. 7a). This assembly consists of a microcontroller that controls a stepper motor whose shaft was fitted with a 3D-printed housing for a permanent magnet (12 mm-diameter nickel-plated neodymium). The north-south axis of the permanent magnet was aligned parallel to the metal disk to ensure effective rotation. The feature could be used on the stand-alone chamber or be integrated into the live-cell imaging process such that the brain tissue could be concurrently rotated while imaged with the confocal microscope (Supplemental Video 3).

**Fig. 7:**
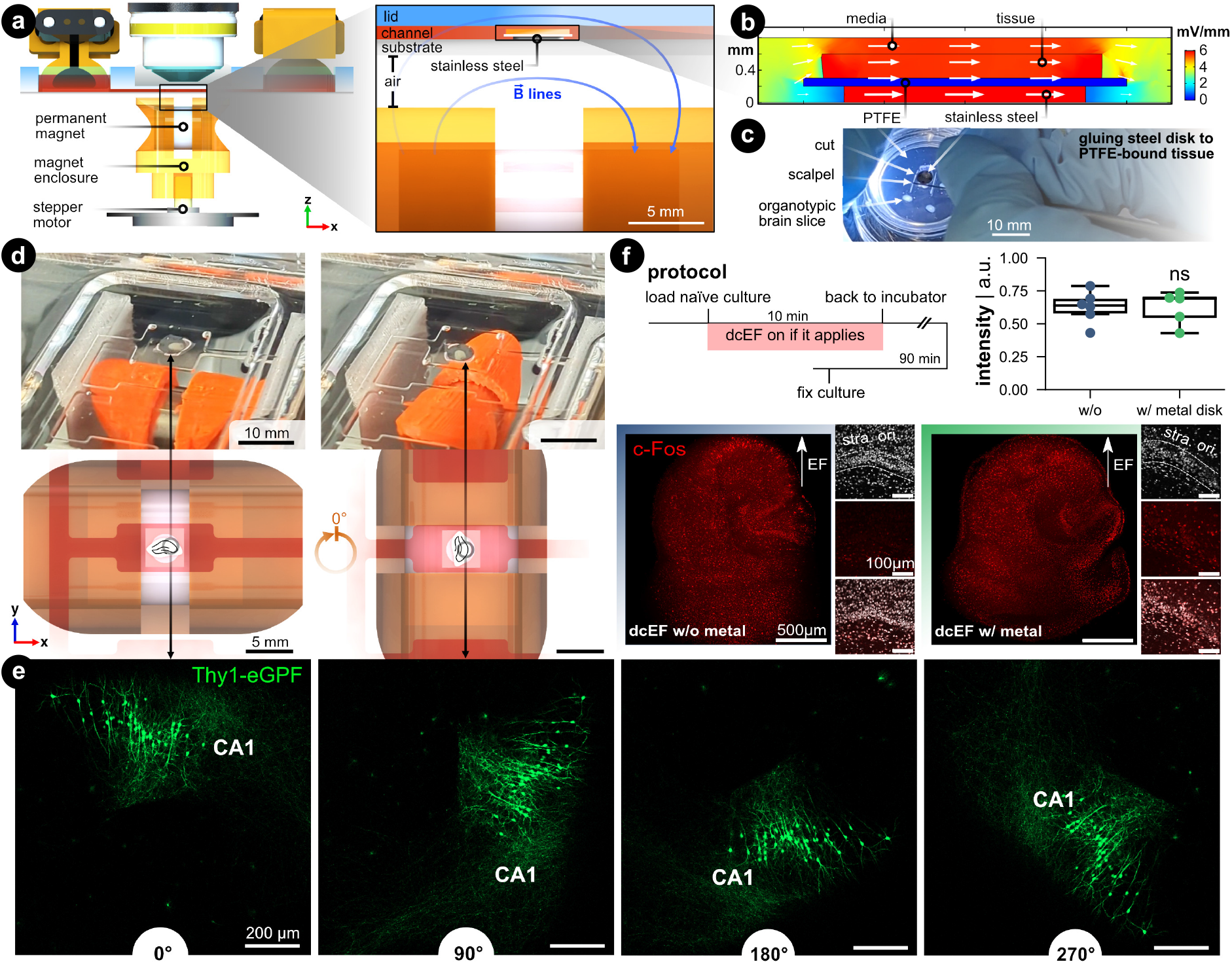
Controlling the brain slice orientation within the microchannel. (a) Side view of the microfluidic stimulation platform with the rotational add-on. The expanded view shows a cross-sectional view of the s1 microchannel and illustrates the added features that enable rotational control: (1) a ferromagnetic stainless steel disk under the brain slice and culturing membrane, and (2) a programmable rotating permanent magnet. The metal disk is resting on the silicone-coated substrate and is not physically attached. The permanent magnet is housed in a 3D-printed enclosure that fits onto a stepper motor shaft whose rotation is controlled with a microcontroller. The magnet’s north-south axis is aligned parallel to the microfluidic chamber, which can be seen with the blue magnetic field lines. (b) FEA simulation of the EF distribution within the s1 microchannel using the same input current used in Fig. 2. Supplemental Fig. S9 expands on this simulation. (c) Image of cutting the tissue/PTFE/disk stack. Note the disk was glued to the underside of the membrane. (d) Demonstration of the *in situ* brain slice rotation within the microchannel. The rendering shows a top view of the rotating brain tissue within the s1 microchannel. (e) Rotation of Thy1-eGFP tissue cultures to further visualize the *in situ* rotation in 90° steps. Note the CA1 region as a datum reference in all rotation images. (f) c-Fos expression after dcEF with and without the metal disk. The insets display the DAPI and c-Fos signal in the CA1 region of the corresponding culture. The box plot shows the c-Fos intensity post-dcEF (4.7 mV mm*^−^*^1^). Mean and quartiles of the whole culture c-Fos signal intensity in two groups (*N* = 6 and *N* = 5 for the dcEF-treated cultures without and with the metal disk, respectively). Mann-Whitney U test was applied for statistical analysis.

FEA simulation was leveraged to validate if the presence of a metal disk in the EF’s path does not affect the EF strength and distribution. First, the DC resistance of the electrolyte-metal interface for the metal disk was measured by electrochemical impedance spectroscopy (EIS) and an equivalent circuit was used to find the charge and mass transfer resistances (Supplemental Fig. S9). The measured ionic conductivity (*σ* = 0.15 S m*^−^*^1^) rather than the metal’s electric conductivity value (*σ* = 1.40 *×* 10^6^ S m*^−^*^1^)

was used as input for the simulation to model the ion-electron current transduction resistance. FEA analysis validated that the EF remained predominately high in the tissue and medium compared to the metal (Fig. 7b). Quantitatively, the EF within the tissue increases a modest 14 % compared to the metal-free analog, which is a consequence of the reduced cross-section (Supplemental Fig. S9d). A protocol was developed that adheres the metal disk to the PTFE-bound tissue (Fig. 7c), to facilitate concurrent rotation of the substrate and tissue within a microfluidic domain Fig. 7d).

To visualize the *in situ* rotational control incorporated into the live-cell imaging process, we placed Thy1-eGFP cultures on the metal disk substrates inside the microfluidic chamber under the microscope. With a preprogrammed button-activated controller which moves the disk in 90 degree steps (Fig. 7d and Supplemental Video 3), we demonstrated the rotational progress within the microchannel (Fig. 7e).

To demonstrate whether the simulated 14 % elevation in EF intensity would experimentally boost the stimulation effects, we stimulated the cultures with the dcEF strength that has been shown not to induce changes in c-Fos expression nor calcium activities (*i.e.*, 4.7 mV mm*^−^*^1^, Supplemental Fig. S7). The extra contribution to dcEF intensity from the presence of a metal disk during stimulation follows suit with the previously observed results by causing upregulation of c-Fos expression (*p* = 0.93, Mann Whitney U test, Fig. 7f) and increasing calcium spikes (Supplemental Fig. S10). The extra handling steps of mounting the cultures onto the metal disk had no influence on cell viability as judged by DAPI signals in the CA1 region (small insets). Also, the metal did not cause any immune responses as verified by analyzing the TNF*α* expression (Supplemental Fig. S11). Our data established that the added rotational design could effectively control the culture orientation inside the microchannel while the internal ferromagnetic disk neither disturbs the EF intensity nor harms culture viability.

## Discussion

Effective and safe use of non-invasive brain stimulation techniques requires addressing questions such as optimal tDCS/tACS electrode montages,^56–58^ magnetic coil orientation,^59^ dose-response relationship,^60^ safety limits of acute and accumulated stimulation intensity,^61^ and stimulation interval’s impact in multisession stimulation.^62^ The answers to these questions have begun to emerge over the past several decades through *in vivo* rodent experiments, computational modeling, and *in vitro* studies using direct or alternating EFs. However, precise control of EF intensity and orientation is paramount to quantitatively link animal experiments with human applications. It is also not trivial to keep the tissue alive over a sufficient experimental time window to allow an in-depth and multi-parametric study. Meanwhile, the prevalent system of stimulating acute slices with parallel electrodes in elaborate electrophysiological setups is not satisfying regarding its limitations in delivering precise dcEF around brain tissue or obtaining results over a relatively long time course.^32^ To address these issues, we developed an inexpensive and high-throughput solution using microfluidic techniques to achieve precise control of uniform EFs in space and time. We displayed various application scenarios using organotypic tissue cultures by extending experiment durations and accommodating cutting-edge probing techniques.

Our microfluidic chamber enables simultaneous live-cell imaging, dcEF stimulation, and modulation of brain slice orientation using an external magnetic rotation assembly. We demonstrated its versatility using mouse tissue cultures, which were monitored for online effects or harvested for offline measurements or further cultivation and examination at later time points. The system’s compact electrode assembly allows for capacitive DC stimulation without the risk of faradaic and potentially toxic reactions or metal ion release. The ability to change the slice orientation during an experiment is time-efficient and avoids potential inter-subject variation while exploring the impact of field orientation, which, next to field strength, is imperative for the outcome. Particularly, the cost of equipment and materials needed for the platform is not prohibitive, and the microfabrication techniques used do not require a cleanroom nor other uncommon infrastructure. Only two commonly-used rapid prototyping equipment are required: CO_2_ laser and plasma chamber. By accurately targeting dcEF intensity, modulating brain slice orientation, and precisely controlling the biochemical microenvironment around the brain tissue, we provide an alternative system to replicate the milestone discoveries in (t)DCS studies and to further bridge *in vitro* studies with human applications.

The design of microfluidic devices and the choice of materials are pivotal for merging advanced functionality with ease of use and parallel experimentation. A microfluidic architecture that generates multiple dcEF strengths from a single pair of electrodes simplifies the setup (*i.e.*, fewer materials, wiring, and connections) and reduces costs (*i.e.*, fewer total devices and constant current sources needed). Furthermore, using open reservoirs allows for easy integration of electrodes that interface with the electrolyte-filled reservoirs. The removable electrodes can easily be scaled in size to match the volume of the reservoir, allowing for longer capacitive DC stimulation times.^50^ In this work, air plasma-treated silicone was reversibly sealed to enable tissue recovery after stimulation. Cell viability tests endorsed the platform’s effectiveness in CO_2_/O_2_ exchange. A completely sealed chamber with inlet and outlet tubing could achieve similar effects via active pressure-driven or peristaltic fluid flow, particularly when substantially long experimental durations are needed. However, such a design would come with increased operational difficulty. It may require a stronger reversible sealing achieved through surface chemistry treatments or coatings. The electrodes should also be integrated within the microfluidic channels or rely on unwieldy salt bridges protruding from the chamber. If long stimulation periods or repetitive stimulation patterns are of great interest, performing the stimulation within incubated microscopes or retrieving cultures for re-culturing back in the regular incubator between sessions might be optional. Furthermore, this microchannel design could be simplified to a single straight rectangular channel to facilitate a lower fluidic/electric resistance so that less potential is needed for a given input current, thus allowing for longer capacitive-dominant stimulation times. Along with the microchannel geometry, the electrode material choice is also imperative in the duration of capacitively-dominant dcEF generation.

An important factor to consider when supplying direct current within an electrolyte is that electrochemical faradaic reactions are inevitable once the electrode’s electrochemical double layer is fully charged/discharged.^42, 63^ Continued faradaic reactions inevitably generate pH shifts, promote tissue oxidative stress via electrochemically-generated ROS, and develop anodal metal ion dissolution that can be toxic to the tissue. To avoid this risk, it is important to carefully consider the electrochemical properties of the electrodes (*e.g.,* charge storage) and the buffering capacity of the medium. Using non-metal electrodes, such as the conducting hydrogel (PEDOT:PSS) used here, can eliminate the concern of toxic metal ion dissolution while providing a material with high ionic capacitance that facilitates constant current discharge over many minutes of DC stimulation. These kinds of new electrode materials show tremendous potential for bridging in-buffer to on-skin uses, which makes them promising for translational applications. Staying within the limits of the capacitive discharge current regime reduces the risk of harmful electrochemical reactions. Although not further explored in this paper, prediction of the capacitive *versus* pseudocapacitive current contributions can be achieved through the Trasatti^64^ or Dunn-generalized Conway method.^65, 66^

We showed that a weak dcEF at 4.7 mV mm*^−^*^1^ failed to trigger c-Fos overexpression, did not increase calcium activity, and unsuccessfully altered synaptic transmission. This suggests that there was no induction of activity-dependent synaptic plasticity at this sub-threshold dcEF intensity. It should be noted here that the vast majority of *in vitro* tDCS studies use Ag/AgCl electrodes without salt bridges to generate similarly weak dcEFs, which are susceptible to faradaic by-product gradients (pH,^37^ Ag^+^,^37, 42^ or ROS^67^) and not just the weak dcEFs. Some studies have shown that a dcEF intensity as low as 0.75 mV mm*^−^*^1^ could trigger long-term potentiation (LTP) like effects^31^ and 4.7 mV mm*^−^*^1^ is within this neural activation range.^16, 46^ According to our recent technical review,^32^ it is highly possible that the actual EF intensities used in these studies were higher than 4.7 mV mm*^−^*^1^, due to a systematic underestimation of dcEF intensity in these studies using conventional calibration approaches.^24–26, 68^ Voroslakos and colleagues have also raised the opinion that the intensity needed for exerting active tDCS effects should be higher than regularly reported literature values.^26^ Yet, Fritsch and colleagues^31^ documented that to induce LTP in their study with a reported 0.75 mV mm*^−^*^1^ dcEF intensity, both the presence of neuromodulator brain-derived neurotrophic factor (BDNF) and a reasonable amount of synaptic activity are required. To achieve this, they applied extra 0.1 Hz pulses to the vertical input pathway. Given that inducing synaptic plasticity is reliant on a plethora of factors, a standardized platform that generates precisely controlled dcEFs and allows for feasible manipulation of neuromodulators would be ideal for solving the puzzling dose-dependency effect for future tDCS studies. Our design definitely serves as a reliable candidate in this regard.

The benefits of the microfluidic chamber for *in vitro* studies extend beyond tDCS and are easily transferable to tACS and rTMS *in vitro* studies by changing the EF type. We showcased this possibility by delivering strong pulsed dcEFs at 140 mV mm*^−^*^1^, which is close to the intensities in rTMS studies.^52, 53^ Despite the high EF intensity, the total charge delivered was much lower for the high current short pulses (1.29 mA *·* 0.1 s *·* 100 pulses = 13 mC) compared to the low current long pulse (43.5 µA *·* 10 min = 26 mC). This means the total delivered charge is also much less than what is ionically stored in the hydrogel and the electrodes likely will not reach faradaic reactions during the high current pulsing. Therefore, our microfluidic chamber and hydrogel electrodes are also compatible with using pulsed dcEFs to mimic rTMS pulses. The controlled microenvironment may also be feasible for accommodating human brain tissue to mimic a miniature “brain” immersed in aCSF and subjected to global electrical stimulation. The integration of non-contact imaging of the microfluidic chamber makes it possible to study neurons, glial cells, and their interaction in the presence of an EF by imaging the calcium signals or cell morphology in real time. Naturally, glial cells participate in brain stimulation-triggered synaptic plasticity,^69, 70^ but conventional approaches are limited in revealing the full story of these cells. These are the kind of questions that we hope the presented platform can address via its versatile live-cell and *post hoc* analysis capabilities. This platform can also be adapted for other explanted tissue types and prove useful for different bioelectronic stimulation applications such as wound healing, morphogenesis, or osseointegration. In summary, the microfluidic chamber inherits the merits of electrotaxis systems and microfluidic techniques to generate precise control of the EF strength and microenvironment. It provides the possibility to verify and build on the cornerstone discoveries of the field. This can be achieved through the platform’s complimentary functionalities with increased throughput, precision, and multimodal measurements. All of which can be integrated with full flexibility in terms of cutting-edge neuroscience methods, particularly enabled by the reversible seal approach and the transparent chamber.

## Materials and Methods

### Simulation of EF using finite element analysis

The microfluidic network, brain slice, and membrane were designed and exported (IGS file extension) in Solidworks (version 2021). The size of the brain slice, which is organotypic entorhinal-hippocampal slice tissue culture in this case, has an area of 5.5 mm^2^ and thickness of 0.30 mm. The polytetrafluoroethylene (PTFE, Teflon) membrane, on which the slice was cultured, was modeled as an area of 16 mm^2^ and thickness of 0.10 mm. COMSOL Multiphysics^®^ software (version 5.3) was used to simulate EF distribution and magnitude using the Electric Currents module. For EF distribution, electrodes sat on top of the reservoirs and were modeled to have an electrical conductivity of PEDOT:PSS hydrogels (*σ* = 2000 S m*^−^*^1^).^71^ The media was modeled after 10 mM phosphate-buffered saline (1*×* PBS) and artificial cerebrospinal fluid (aCSF), both of which have an electrical conductivity (*σ*) of 1.5 S m*^−^*^1^.^72^ This was also measured/verified using a portable conductivity meter (DiST6 EC/TDS, Hanna Instruments, Germany). The conductivity used for tissue and PTFE membrane was 0.5 and 1 *×* 10*^−^*^16^ S m*^−^*^1^, respectively.^73, 74^

The relative permittivity is 5 *×* 10^7^, 2, and 80 for tissue,^73^ PTFE,^74^ and media, respectively. The cathode was set to 0 V. The PTFE membrane was modeled as an ideal insulator due to the 15-orders of magnitude difference to the next closest material in the system. The input current density (placed at the anode face) was swept in order to identify which input current is needed to achieve EF strengths around 1 mV mm*^−^*^1^ in the tissue-containing reservoirs. In the case including the stainless steel disk, all the same conditions were used and are described in more detail within Supplemental Fig S9.

### Preparation of microfluidic devices

For a process workflow with images, please see Supplemental Fig. S4a-d. Poly(methyl methacrylate) (PMMA, acrylic) sheets were cut using CO_2_ laser (Beambox Pro, Flux, Taiwan). This acrylic (Modulor, Germany) mold was made of five distinct components: substrate (3 mm thick), fluidic negative (0.8 mm thick), reservoir negatives (8 mm thick), side walls (8 mm thick), and immersion objective trough (8 mm thick). The first four components were solvent-bonded together using dichloromethane. A freshly mixed two-part silicone (Sylgard 184, Dow Corning, USA) called polydimethylsiloxane (PDMS) was poured into the acrylic mold. This PDMS was filled until it covered the microchannels and degassed for 30 min in a vacuum desiccator, then it was cured at 70 °C for 1 h. The final acrylic piece was then placed over the cured microchannels and more PDMS was poured over the mold to further define the reservoirs and trough for the immersion microscope objective, then finally cured for 1 h at 70 °C. Meanwhile, PDMS was also poured into one-well polystyrene plates (Cellstar, Greiner Bio-one, Germany) and cured. On the day of experiments with brain slices, both the molded PDMS device and PDMS substrate (*i.e.,* one-well plate) were air plasma-treated for 30 W for 30 s (Femto Model 1B1, Diener Electronic, Germany) to increase the hydrophilicity of the naturally hydrophobic PDMS (by oxidizing the surface to create more silanol (Si-O-H) and hydroxy (C-O-H) groups to allow better fluid flow and to improve temporary, reversible bonding between PDMS layers. The low power plasma treatment, the subsequent hydrophobic recovery (high surface energy reconfiguring to a lower energy state), and the smoothness of the PDMS all play a role in the strength of the bond between PDMS layers. In other words, if too high of power is used or the two treated surfaces adhere too quickly after exposure, then the bond might be too strong to recover the tissue after sealing. Contrarily, if not enough power is used, then PDMS might not be hydrophilic enough for easy fluidic loading and the bond might not be strong enough to endure simple hydrostatic flow. We found that devices had functioned optimally within 1 to 4 h after the aforementioned plasma treatment settings.

### Preparation of electrodes

Laser-induced graphene (LIG) was made with a mid-IR (wavelength of 10.6 µm) CO_2_ laser (VLS 2.30, Universal Laser Systems, USA) by carbonization of a 75 µm-thick polyimide (PI) sheet (Kapton HN, Dupont, USA).^43^ In order to improve electrical conductivity and ability to store ions (*i.e.,* electrochemical charge storage capacity), a conducting hydrogel (poly(3,4-ethylenedioxythiophene) polystyrene sulfonate - PEDOT:PSS) was coated on the LIG. In short, the PEDOT:PSS dispersion (1.3 % in water) was spiked with 15 % dimethyl sulfoxide (DMSO) and cast onto the amine-functionalized and polyurethane-coated LIG, which follows our previously described work.^43^ Electrode connection lines (*i.e.,* between electrical bump pad and electroactive area) were insulated by coating with an acrylate-based varnish (Essence 2 in 1, Cosnova). PEDOT:PSS hydrogel coated LIG electrodes were stored in 1*×* phosphate-buffered saline (PBS) until further use. For Fig. 6, the electrodes were doubled. Two electrodes were placed back-to-back and coated with a 1*x* PBS-infused agarose to buffer any potential electrochemical by-products (see Supplemental Fig. S8 for more detail).

### Preparation of the rotational control of brain slice

The programmable rotating magnet was made with four components: a microcontroller (Arduino, Uno SMD R3, Italy), a stepper motor (28BYJ48, Adafruit Industries), a 3D-printed (PET-G, Prusa i3 MK3S, Czech Republic) magnet holder that fits into the stepper motor shaft, and a 12 mm-diameter and 16 mm-long nickel-plated neodymium permanent magnet. The microcontroller was programmed using Arduino IDE to communicate with a three-button controller where each button tells the stepper motor to either rotate in 45°-steps clockwise, or 90°-steps clockwise or counterclockwise. Another added 3D-printed part was made to incorporate into the microscope stand so that the magnetic assembly could hang below the microchamber. This entire assembly is visualized in Supplemental Video 3.

Stainless steel sheets (1.4310 grade, 200 µm-thick) were cut with a near-IR (wavelength of 1.064 µm) laser (DPL Genesis Marker Nd:YAG, ACI Laser GmbH, Germany). The laser settings used was a power of 4.5 W (*i.e.,* 100 %), velocity of 1.0 mm*/*s, frequency of 500 Hz, pulse width of 3.0 µ sec, and 20 overall passes. Subsequently, the 4.0 mm metal disks were sonicated for 10 min in 1 % acetic acid in 70 % ethanol to get rid of any oxidized steel from the lasing process. The disks were stored in a sterile container until further use.

For adhering to tissue cultures, a cyanoacrylate-based glue (Histoacryl Blue, Braun Surgical, Spain) that is common for histological slicing was used to glue the metal disk to the PTFE membrane (Supplemental Video 3). The PTFE insert/membrane that contains the organotypic slides was removed from the well-plate and flipped upside-down so that the membrane was facing up. The metal disk was placed onto the membrane and directly above the target slice. A de-insulated copper wire was used as the glue applicator by dipping into a vial of the uncured blue glue and dabbing the wet wire tip around the edge of the metal disk. Note that the glue cures once wetted by the media present on the membrane. The entire construct (disk/insert/slice) is now flipped over and a scalpel is used to cut out a roughly 4 mm by 4 mm square. The rest of the chamber assembly process follows suit with the *Chamber exposure and dcEF stimulation* section below.

### Ethics statement

In order to justify the safety of the silicone dcEF microfluidic chamber, we used organotypic entorhinal-hippocampal tissue cultures prepared from mouse pups at 3 to 5 days post-birth (P3-P5) from different mouse lines. All animals were kept under a 12 h light-dark cycle with food and water provided *adlibitum*. One male and one or two female(s) were kept within the same cage for mating. All experiments were performed according to German animal welfare legislation and approved by the appropriate animal welfare committee and the animal welfare officer of Albert-Ludwigs-Universität Freiburg, Faculty of Medicine under X-17/07K, X-21/01B, X-17/09C, and X-18/02C. All effort was made to reduce the pain or distress of animals.

### Preparation of tissue cultures

All tissue cultures were prepared at P3-P5 from C57BL/6J, C57BL/6-Tg(TNF*α*-eGFP),^51, 75^ and Thy1- eGFP mice of either sex as previously described.^76^ Incubation medium (pH = 7.38) contained 50% (v/v) minimum essential media (#21575 *−* 022, Thermo Fisher, USA), 25% (v/v) basal medium eagle (#41010 *−* 026, Gibco, Thermo Fisher, USA), 25% (v/v) heat-inactivated normal horse serum (#26050 *−* 088, Gibco, Thermo Fisher, New Zealand), 25 mM HEPES buffer solution (#15630 *−* 056, Gibco, Thermo Fisher, USA), 0.15% (w/v) sodium bicarbonate (#25080 *−* 060, Gibco, UK), 0.65% (w/v) glucose (G8769- 100ML, Sigma-Aldrich, UK), 0.1 mg*/*mL streptomycin, 100 U*/*mL penicillin, and 2 mM GlutaMAX (#35050 *−* 061, Gibco, China). All tissue cultures were cultured for at least 18 days inside the incubator with a humidified atmosphere (5% CO_2_ at 35 °C. The incubation medium was renewed every Monday, Wednesday, and Friday.

### Chamber exposure and dcEF stimulation

For a process workflow with images, please see Supplemental Fig. S4e-h. In order to load individual cultures into the chamber, we first readied a plasma-treated silicone-coated one-well plate and added an acrylic stencil with rectangular cut-outs that matched the size and location of the stimulation zones of the molded microfluidic device. Next, we cut through the PTFE membrane where the slices have been growing and quickly transferred (*≈* 10 s) the slice with membrane onto the stimulation zone of choice (guided by the aforementioned stencil). The stencil was immediately removed. Then, we encapsulated the wet tissue slice(s) with the de-molded and plasma-treated silicone microfluidic chamber to completely define the microfluidic environment. Immediately after sealing, 5 mL of incubation medium was injected into the inlet/anode reservoir. The medium flows into the three channels and fills the other reservoir via capillary and hydrostatic pressure forces. If not otherwise stated, we used the same fresh incubation medium that was used for culturing the cultures inside the incubator, and it was always pre-warmed to 35 °C and pH adjusted to pH = 7.38. For the chamber exposure group, cultures were randomly placed into the three channels and remained immersed. In the dcEF stimulation experiment, the cultures remained immersed in the corresponding channels for certain EF intensities with the same orientation across all trials. The whole plate was seated on the live-cell microscope stage that was constantly held at 35 °C and remained there throughout the experiment. For the dcEF stimulation group, two PEDOT:PSS hydrogel electrodes were immersed in the two reservoirs and connected to a constant current source (PGSTAT101, Metrohm Autolab, Switzerland). For weak dcEFs, the input current was 43.5 µA, while for the strong pulsed dcEFs the input current was pulsed for 0.1 s at 1.29 mA with 0.5 s rest periods for a total of 100 pulses. Imaging experiments, if planned, were performed when the cultures were housed inside the chamber. After chamber exposure or dcEF stimulation, the silicone cover was peeled to retrieve the cultures. The cultures were either subject to offline measurements or placed back onto the incubation PTFE insert and re-cultured in the incubator for later examination.

### SYTOX-green staining

In order to first examine whether the microfluidic chamber affects cell viability, we used tissue cultures prepared from wild-type animals and stained the nuclei of dead cells with NucGreen Dead 488 (SYTOX-green #R37109, Thermo Fisher, USA). Tissue cultures were submerged inside the incubation medium within the microfluidic chamber for 20 min. After 15 min of incubation, 250 *µ*L SYTOX-green was added into the open inlet reservoir to incubate the cultures for another 5 min. In order to further examine the potential cell death caused by dcEF stimulation inside the chamber, we used the same procedure but now with the constant current source turned on during the last 10 min, including the last 5 min of SYTOX-green incubation. For näıve controls, the cultures remained inside the incubator the whole time and incubated with 250 *µ*L SYTOX-green spiked into 5 mL incubation medium for the last 5 min. For positive controls, cultures were treated with 50 *µ*M N-methyl D-aspartate (NMDA) for four hours in an interface manner inside 1 mL incubation medium and later incubated for SYTOX-green as described for näıve controls. After 5 min of incubation time with SYTOX-green, all the cultures were fixed for DAPI staining and confocal microscope imaging.

### Live-cell microscope imaging

The live-cell microscope was used in three experiments, (1) to inspect whether the microfluidic chamber irritates tissue cultures and causes potential inflammatory status by examining the expression of TNF*α*, (2) to monitor the calcium dynamics of the CA1 region via calcium imaging, and (3) to monitor the rotational control of culture orientation within the microchannel. Live-cell imaging was performed at a Zeiss LSM800 microscope with 10*×* water-immersion objective (W N-Achroplan 10*×*/0.3 M27; #420947- 9900-000; Carl Zeiss). To image the culture inside the microfluidic chamber, we immersed the objective into a pre-designed well on top of the chamber filled with distilled water. Note that the microchannel’s silicone lid separates the culture media-filled microchannel and the water-filled immersion well. In some conditions, we imaged the näıve cultures without the microfluidic chambers, but rather inside a 35 mm Petri dish as described before.^70^ Then the filter insert with 2 to 4 cultures was placed inside the Petri dish containing 5 mL pre-warmed pH-adjusted incubation medium.

### Live-cell monitoring of TNF***α*** expression

The expression of TNF*α* was conducted with tissue cultures prepared from TNF*α*-reporter animals C57BL/6-Tg(TNF*α*-eGFP).^51^ For the chamber immersion group, the cultures were imaged immediately after being loaded into the dcEF microfluidic chamber and imaged again after 20 min’s incubation. For the dcEF stimulation group, the same procedure was applied while the current source was switched on to apply dcEF stimulation for the last 10 min and the first 10 min being incubation and operation time. For positive controls, tissue cultures were treated with bacterial lipopolysaccharide (LPS from Escherichia coli O111:B4, #L4391, Sigma-Aldrich, UK, 1 *µ*g*/*mL) for 3 days inside the incubator and imaged before and after 3 days of treatment as previously described.^51^

### Calcium imaging

Calcium imaging was conducted in tissue cultures prepared from wild-type animals. The entire tissue culture (between DIV3 and DIV5) was transfected with 1 *µ*L AAV1-hSyn1-GCaMP6f-P2A-nls-dTomato virus (Addgene viral prep #51085-AAV1, http://n2t.net/addgene:51085, RRID:Addgene 51085, a gift from Jonathan Ting) diluted 1 : 4 in 1*×* PBS (pH = 7.38), by pipetting a drop of the mixture on top of the culture to cover the entire culture. The virus will express calcium indicator GCaMP6f as well as TdTomato in transfected neurons. To examine if handling and incubation within the chamber will affect neural activity level, we used the within-subject experimental design. Näıve control cultures were first imaged within a 35 mm Petri dish and then loaded into the microfluidic chamber and imaged after 10 min. These cultures were later on split into two groups to examine the impact of a weak dcEF stimulation (see Supplemental Materials for details). The positive controls were also imaged twice for two consecutive sessions. During the first session, cultures were imaged immediately after being loaded into the chamber; then we turned on pulsed dcEF stimulation with high intensity and imaged the cultures for another session. To cross-validate the results of whole-cell patch-clamp recording, we focused on the CA1 area as our region of interest (ROI) for calcium imaging. Videos were captured at 128 px *×* 128 px resolution for 2 min with an interval of 120 ms.

### Imaging Thy1-eGFP cultures

Thy1-eGFP cultures were imaged with a live-cell microscope to visualize the real-time *in situ* rotational control. Images were taken after each rotation.

### Whole-cell patch-clamp recording

Whole-cell patch-clamp recordings were carried out to investigate the synaptic transmission function and neural intrinsic membrane properties. Four groups were designed. Näıve controls came directly from the incubator. Chamber-only cultures were recorded immediately after being immersed inside the microfluidic chamber for 20 min. After the dcEF stimulated cultures, they were immersed inside the chamber for a total duration of 20 min, during which a weak dcEF was delivered for 10 min in between. Since altered synaptic transmission was commonly regarded as a readout of synaptic plasticity, which takes time to occur, we retrieved the stimulated cultures and re-cultured them back into the regular incubator for 2 *±* 1 h before recording to allow for plasticity induction, if there is any. Therefore to rule out the confounding roles of retrieving and re-culturing, we prepared and recorded another set of chamber control cultures when through the same retrieving and re-culturing process as dcEF-stimulated cultures. The bath solution aCSF consists of (in mM) 126 NaCl, 2.5 KCl, 26 NaHCO_3_, 1.26 NaH_2_PO_4_, 2 CaCl_2_, 2 MgCl_2_, and 10 glucose. It was continuously oxygenated with 5% CO_2_/95% O_2_ and warmed up to 35 °C. The internal solution for patch pipettes contained (in mM) 126 K-gluconate, 10 HEPES, 4 KCl, 4 ATP-Mg, 0.3 GTP-Na_2_, 10 PO-Creatine, 0.3% (w/v) biocytin (pH = 7.25 with KOH, 290 mOsm with sucrose). The patch pipettes have a tip resistance of 4 MΩ to 6 MΩ. CA1 pyramidal neurons were identified using a LN-Scope (Luigs & Neumann, Germany) equipped with an infrared dot-contrast 40*×* water-immersion objective (NA 0.8; Olympus). For synaptic transmission function, spontaneous excitatory postsynaptic currents (sEPSCs) were recorded in a voltage-clamp mode at a holding potential of *−*70 mV. Series resistance was monitored before and after each recording. Intrinsic membrane properties were recorded in the current-clamp mode, where pipette capacitance of 2 pF was corrected and series resistance was compensated using the automated bridge balance tool of the MultiClamp commander. I-V-curves were generated by injecting 1 s square pulse currents starting at *−*100 pA and increasing up to 500 pA for every 10 pA. The sweep duration is 2 s.

### Tissue fixation and immunohistochemical staining

In some experiments, *post hoc* confocal microscope imaging was applied after tissue fixation and immuno- histochemical staining. These cultures were fixed by immersing into cold 4% (w/v) paraformaldehyde (PFA) in 1*×* PBS with 4% (w/v) sucrose for 1 h and transferred into 1*×* PBS for storage at 4 °C after being washed in 1*×* PBS. Later, we did fluorescent staining on fixed cultures to visualize neurons or proteins of interest.

#### Streptavidin staining

Streptavidin staining was used to visualize the CA1 pyramidal neurons recorded during whole-cell patch-clamp recording. All recorded and fixed cultures were first washed three times with 1*×* PBS (3 *×* 10 min) to remove residual PFA. We then incubated the cultures with Streptavidin 488 (1 : 1000, #S32354, Invitrogen, Thermo Fisher, USA) in 1*×* PBS with 10% (v/v) in normal goat serum and 0.05% (v/v) Triton X-100 at 4 °C overnight. In the next morning, cultures were rinsed with 1*×* PBS (3 *×* 10 min) and incubated with DAPI (1 : 2000) for 20 min. After another 4 washes with 1*×* PBS (4 *×* 10 min), we mounted the cultures on glass slides with DAKO anti-fading mounting medium (#S302380 *−* 2, Agilent) for confocal microscope imaging.

#### c-Fos staining

In order to probe the neural activation effects of dcEF in the presence of a meral dick or not, we also stained cultures with immediate early gene c-Fos. We treated the cultures for 10 min with 4.7 mV mm*^−^*^1^ intensity. 10 min later, cultures were transferred onto an insert and returned to the incubator for another 90 min to allow for c-Fos expression and then fixed with PFA. A similar procedure was applied to compare the neural activation level between näıve controls, chamber-immersed cultures, and dcEF-stimulated cultures, please refer to the Supplemental Materials for details.

As a means to stain c-Fos, we washed all the cultures three times with 1*×* PBS to remove PFA and then blocked at room temperature (RT) for 1 h with 10% (v/v) in normal goat serum in PBS with 0.5% (v/v) Triton X-100 to reduce nonspecific staining while increasing antibody penetration. Later the cultures were incubated at 4 °C with rabbit anti-cFos (Cat# 226 008, RRID:AB 2891278, Synaptic Systems, 1 : 1000) in PBS with 10% (v/v) normal goat serum and 0.05% (v/v) Triton X-100 for 48 h. Cultures were rinsed with 1*×* PBS (3 *×* 10 min) and incubated again with Alexa568 anti-rabbit (1 : 1000) in PBS with 10% (v/v) normal goat serum and 0.05% (v/v) Triton X-100 overnight at 4 °C. The same washing, DAPI staining, and mounting procedures were performed as for streptavidin staining.

#### DAPI staining for SYTOX-green stained cultures

For cultures stained for SYTOX-green and fixed afterward, we also stained these cultures for DAPI following the previously described procedure.

### Confocal microscope imaging

Leica SP8 laser-scanning microscope was used to acquire fluorescent images of c-Fos, SYTOX-green, and biocytin-filled neurons. We used the 20*×* multi-immersion (NA 0.75; Leica) objective to tile-scan the whole culture stack at 512 px *×* 512 px resolution with a step size of Δ*z* = 2 *µ*m. Laser intensity was adjusted accordingly to achieve comparable non-saturated fluorescence intensity among all groups.

### Quantification and data analysis

#### Quantifying microscope images

SYTOX-green, TNF*α*, and c-Fos signals were quantified by fluorescence intensity with Fiji ImageJ. For the SYTOX-green signal, *z*-stacked images were obtained with maximum projection and raw signal intensity was quantified for the whole culture. For the TNF*α* signal, *z*-stacked images were obtained with maximum projection, and an oval-shaped region of interest (ROI) at 1000 pixels *×* 1000 pixels was drawn to include the subregions of the dentate gyrus (DG), CA3, CA1, and part of the entorhinal cortex (EC). The same ROI was applied to each culture imaged before and after manipulation to quantify the raw signal intensity. For the c-Fos signals, *z*-stacked images of the middle 10 planes were obtained with maximum projection and an oval-shaped region of interest (ROI) at 1000 px *×* 1000 px was applied to cover the whole area of the dentate gyrus (DG), CA3, CA2, and CA1.

#### Quantifying electrophysiological recording data

Excitatory postsynaptic currents were analyzed using the automated event detection tool from the pClamp11 software package as previously described.^77^

#### Analyzing calcium imaging data

The GFP signal intensity of calcium indicator GCaMP6f was quantified frame by frame to extract the time series of individual neurons in CA1. Computer vision algorithms were applied to identify individual neurons sampled in each imaging session. For cell detection, we first averaged over the whole time series data from the tdTomato signal to obtain an average intensity. We used tdTomato for cell detection instead of EGFP because with EGPF some of the dendrites also get detected as cell bodies. We then applied a median blur to remove noise. A morphological opening with a kernel size of 32 was performed to obtain background, which was then removed from the time-averaged frame. We then applied a second morphological opening with a kernel of size 2 to sharpen the cell contours. Next, we performed thresholding based on a value that we found by looking through raw recordings (images at different processing stages could be found in Supplemental Fig. S12). In this work, we used a threshold value of 100. The final step was to detect useful cells and their contours which we obtained by detecting connected components in the image. From these connected components, we only choose cells with a cell area larger than 20 pixels. The final time series data was obtained by a spatial average over all pixels. Raw trace was detrended by subtracting the median value with a rolling method (window size: 20) and then normalized based on the mean of tread in each trace to achieve Δ*F/F*_0_ traces. Calcium spikes were automatically detected for each processed trace with a threshold of 3 times the standard deviation from the mean values. Individual traces and spike detection data were visually inspected for quality control.

#### Statistical analysis

For SYTOX-green and the parameters extracted from whole-cell patch-clamp recordings, The Kruskal-Wallis test followed by Dunn’s multiple comparisons test was used for group-level examination and pair- wise comparisons. For c-Fos signals, the Mann-Whitney U test was applied to compare the data obtained with and with a metal disk. For input-output curve analysis, RM two-way ANOVA followed by Sidak’s multiple comparison tests was used. For the TNF*α* signal, the Wilcoxon test was applied to examine the difference between pre- and post-measurements of individual cultures in three groups. Although the same culture was imaged twice for comparing their calcium activity, the neural identification algorithm did not always return the same population size for extracting activity for individual neurons. Therefore, an unpaired student’s t-test was applied, and the conclusions were double-checked with linear mixed models to rule out the impact of data clustering per culture.

## Supporting information

Video 1 = fluidic and electrical analog

Video 2 = calcium imaging example with a strong pulsed dcEF

Video 3 = magnetic rotation assembly description and demonstration

## Supportive Information

### Funding

This project was funded by the European Research Council (ERC) under the European Union’s Horizon 2020 Research and Innovation program under the grant agreement (No. 759655, SPEEDER). This work furthermore received support from BrainLinks-BrainTools, Cluster of Excellence funded by the German Research Foundation (DFG, EXC 1086), currently funded by the Federal Ministry of Economics, Science and Arts of Baden Württemberg within the sustainability program for projects of the excellence initiative. Animal experiments were funded by the National Institutes of Health, USA (NIH; 1R01NS109498) and by the Federal Ministry of Education and Research, Germany (BMBF, 01GQ1804A).

## Acknowledgements

We thank Monika Paetzold and Susanna Glaser for genotyping the animals and thank Birgit Egle for the lab service. We also thank Lukas Matter and Christian Boehler for fruitful discussions about the magnetic rotation assembly and electrochemical experiments.

## Author contributions statement

A.V. and M.A. conceived the project. S.S. designed and fabricated the microfluidic device, electrodes, and magnetic disk assembly. S.S. performed FEA analysis and electrode characterization. H.L. designed the animal experiments. H.L. and S.S. performed chamber treatment and live-cell imaging. H.L. conducted immunohistochemistry and analyzed microscope imaging data. S.G. extracted calcium traces for individual neurons. M.L. performed electrophysiological recordings. M.L. and H.L. analyzed electrophysiological data. S.S. and H.L. made the figures and co-wrote the manuscript. All authors reviewed the manuscript.

## Additional information

**Accession codes**: All raw data and analysis scripts are available here https://github.com/ErbB4/dcEF-chamber.

