## Supplemental Figures for "On-chip brain slice stimulation: precise control of electric fields and tissue orientation"

**Abbreviated title: On-chip brain slice stimulation**

Sebastian Shaner[1,2*], Han Lu[2,3*], Maximilian Lenz[3], Shreyash Garg[4], Andreas Vlachos[2,3,5,#], Maria Asplund[1,2,6,7,8,#]

[1]Department of Microsystems Engineering, University of Freiburg, Freiburg, Germany

[2]BrainLinks-BrainTools Center, University of Freiburg, Freiburg, Germany

[3]Department of Neuroanatomy, Institute of Anatomy and Cell Biology, Faculty of Medicine, University of Freiburg, Freiburg, Germany

[4]MSc Neuroscience Program, Faculty of Biology, University of Freiburg, Freiburg, Germany

[5]Center for Basics in Neuromodulation (NeuroModulBasics), Faculty of Medicine, University of Freiburg, Freiburg, Germany

[6]Freiburg Institute for Advanced Studies (FRIAS), University of Freiburg, Freiburg, Germany

[7]Division of Nursing and Medical Technology, Luleå University of Technology, Luleå, Sweden

[8]Department of Microtechnology and Nanoscience, Chalmers University of Technology, Gothenburg, Sweden

[*]joint first authorship

[#]joint senior authorship

**Declarations of interest:** none

### Supplementary Notes & Figures

**Note S1: Estimating the capacitive current time based off of slow scan rate cyclic voltammetry titration of supercapacitive PEDOT:PSS hydrogel**

One way to estimate capacitive current time for direct current stimulation is via the common electrochemistry technique of cyclic voltammetry (CV). Simply put, a CV is a dynamic electrochemical method where the potential of the electrode of interest (i.e., working electrode in a three-electrode setup) is ramped linearly versus time and back down to the initial set point to complete one cycle, all of which is taking place in an electrolyte of choice. In order to allow full discharge of the large (15 mm-diameter) and thick conducting hydrogel ($\approx$ 50 - 100 µm), slow CV scan rates (1 - 8 mV s$^{-1}$) are performed to minimize the influence of an ohmic drop. CVs are executed between two voltage bounds before water electrolysis occurs window (i.e., water-splitting window *versus* a Ag/AgCl reference electrode) to see where obvious faradaic currents occur (i.e., a large increase in current's slope). For example, in 1$\times$ PBS a voltage window between $-0.6$ and 0.9 V (versus Ag/AgCl) is used since the faradaic current begins within these distal voltage endpoints for PEDOT:PSS-coated electrodes. If one focuses on the anodic current where this inflection point begins ($\approx$ 0.6 V), then these currents can be plotted versus scan rates (Fig. S2). For a capacitor, the charge ($Q$) is equal to capacitance ($C$) times voltage ($U$). Where $C$ can be calculated from the current ($i$) times the time derivative of $U$, or scan rate ($\mathrm{d}U/\mathrm{d}t$). Therefore, the slope of the $i$ vs. $\mathrm{d}U/\mathrm{d}t$ yields $C$. Assuming strictly faradaic current begins at 0.6 V for these materials in this electrolyte, then $Q = 32.8$ mC. Dividing $Q$ by the constant input current used in experiments (43.5 µA) yields a capacitive current time of approximately 12.5 min.

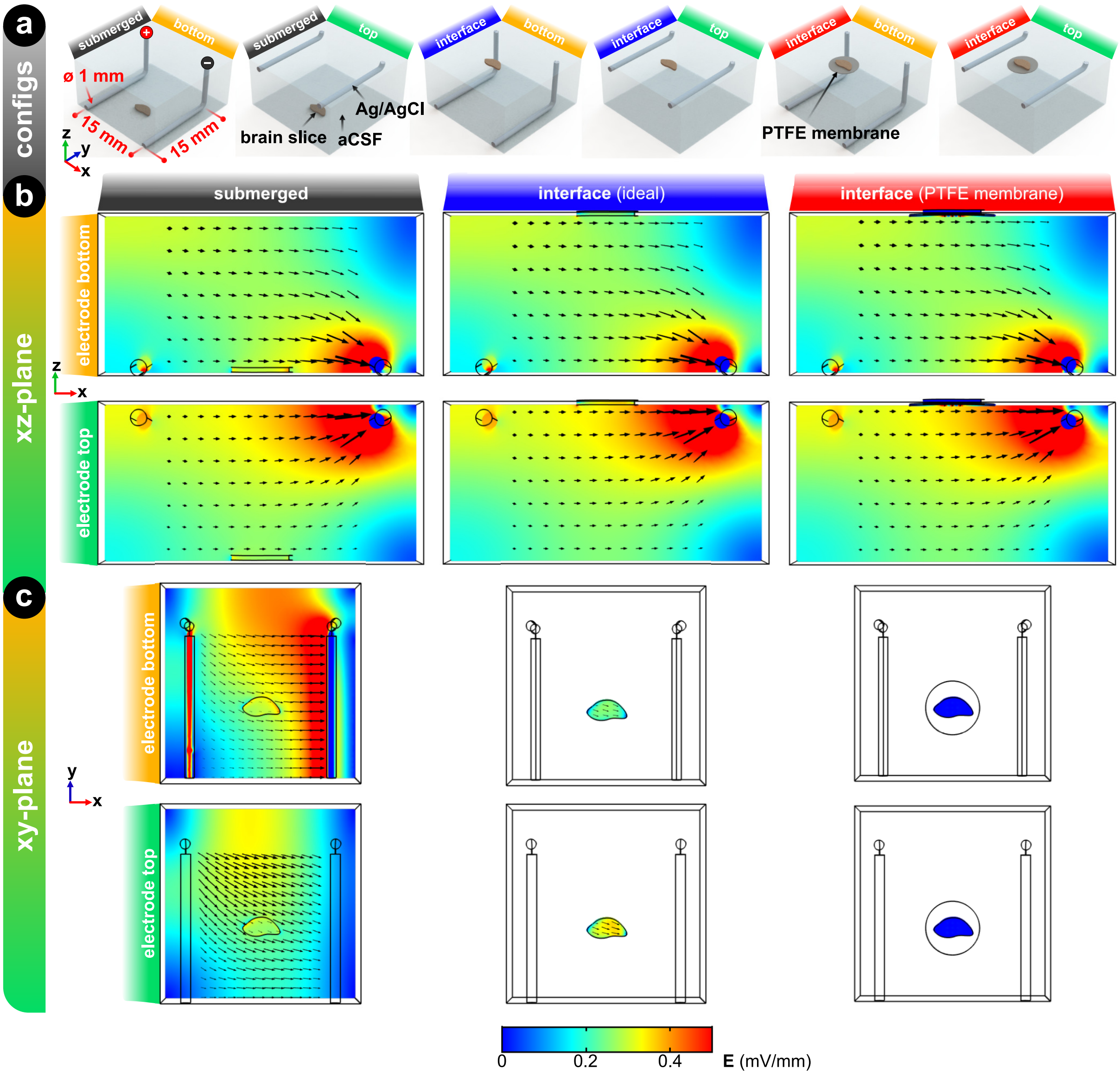


**Fig. S1: Simulation of the EF distribution generated by parallel stimulation electrodes with different orientations relative to the brain slice.** (a) Three-dimensional illustration of six different orientation combinations: a brain slice submerged in aCSF, a brain slice in an ideal "interface" scenario where it sits on top of the aCSF with no insulating support, and an organotypic brain slice with PTFE membrane that has 9 tissue-filled micropores (10 µm diameter) that directly touch the electrolyte, thus mimicking a more realistic "interface" scenario. All of which has electrodes either on the bottom or the top of the aCSF. Electrode diameter (1 mm diameter), length (15 mm), gap (15 mm), and input current (100 µA) were obtained from Bikson et al.[1] and Kronberg et al.[2] Note that for most publications, the z-height of the aCSF/electrolyte is not reported, so we used 10 mm as a reference value. Electrical conductivity values were leveraged from prior publications.[3–7] (b) EF distribution of the mid $xz$-plane of the brain slice between the parallel electrodes for all six cases. (c) EF distribution of the mid $xy$-plane of the brain slice between the parallel electrodes. The EF scale is the same for all, which can be seen at the bottom of the figure.

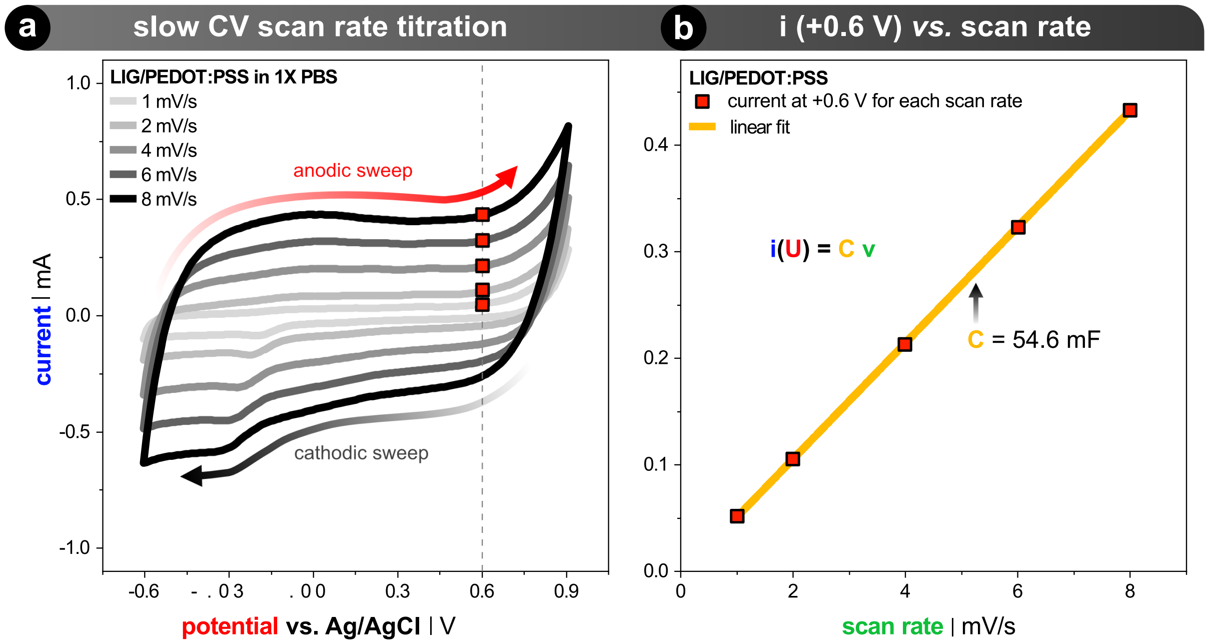


**Fig. S2: Cyclic voltammetry (CV) of PEDOT:PSS hydrogel-coated laser-induced graphene in $1\times$ phosphate-buffered saline at pH 7.4 and room temperature**. (a) CV in a 3-electrode setup (working electrode = LIG+PEDOT:PSS, counter electrode = large stainless steel sheet, reference electrode = Ag/AgCl in 3 M NaCl). CV performed at 5 different slow scan rates ($1$ - $8\,\mathrm{mV\,s^{-1}}$). The anodic sweep voltage of 0.6 V was determined to be the inflection point current starts to turn predominately faradaic. (b) The current values at 0.6 V for each scan rate is plotted and fitted with a linear fit. See Supplemental Note 1 for more information.

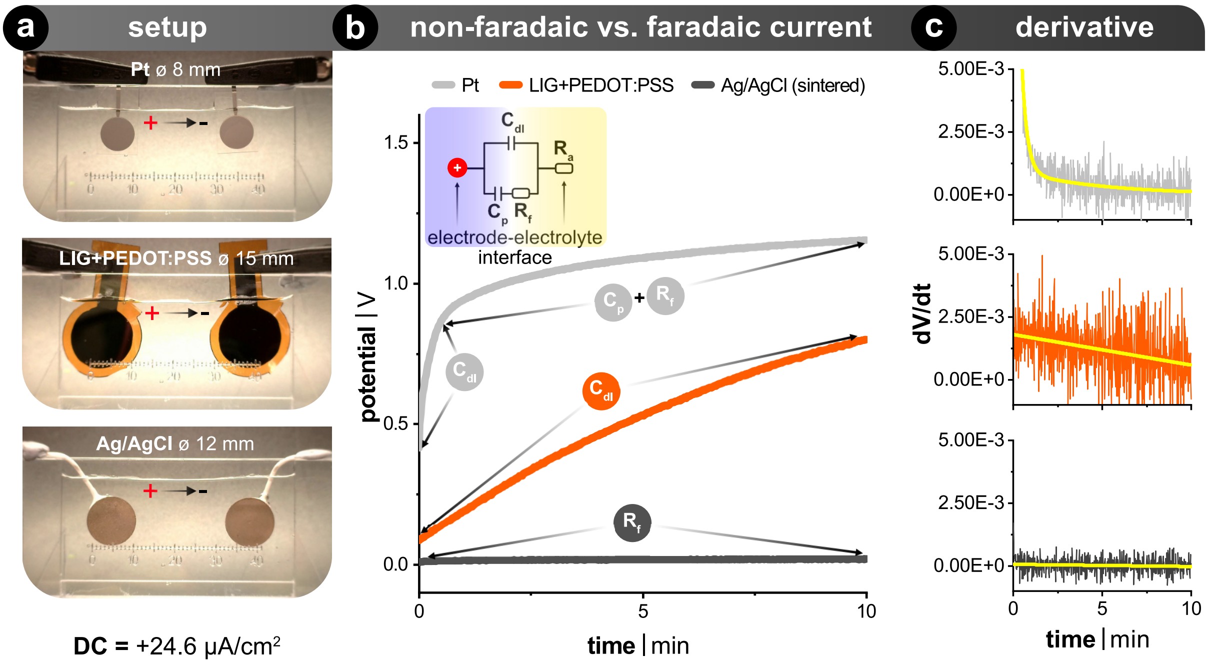


**Fig. S3: Comparison of common metal DC electrode materials versus the non-metal hydrogel electrodes used in this paper**. (a) Example images of the different electrode materials in an acrylic stimulation chamber, which holds 5 mL of 1× PBS. The scale bar that is etched into the chamber is in mm (0–40 mm). The Pt film (<1 µm-thick) was cleanroom fabricated via sputtering onto a glass substrate and insulated with SU-8, the LIG+PEDOT:PSS hydrogel is relatively thin at around 0.1 mm-thick, and 1.0 mm-thick sintered Ag/AgCl electrodes were purchased (Science-Products GmbH, E-204). (b) Voltage excursion plots for the most common (t)DCS electrode material (Ag/AgCl in the superior sintered form), another common metal electrode used for DC (Pt: platinum), and LIG+PEDOT:PSS. Constant current was applied in a two-electrode setup, where the input current density for all cases was $+24.6\,\mu$A cm$^{-2}$. The DC stimulation time of 10 min correlates to stimulation time used throughout in the main text. The upper-left inset shows an equivalent circuit of the electrode-electrolyte interface for the anode, where $C_{\mathrm{dl}}$ is capacitive electrochemical double layer, $R_{\mathrm{f}}$ denotes faradaic charge transfer, and $Z_{\mathrm{d}}$ represents pseudocapacitance that are all in parallel. The electrolyte resistance (i.e., access resistance, $R_{\mathrm{a}}$) is the aqueous connection between anode and cathode. In the potential versus time plot, the initial rising slope is principally due to the $C_{\mathrm{dl}}$. The flattening of this curve is when the faradaic processes $R_f$ become dominant. The transition from predominately capacitive to faradaic current can transition via pseudocapacitive mechanisms, which is material and electrolyte dependant. (c) These are the first derivative of the plots in (b), which helps illustrate if there is predominately capacitive or faradaic reactions occurring, where faradaic currents are easily noticeable when the derivative slope becomes zero. Note that the Pt electrode has a rapid $C_{\mathrm{dl}}$ discharge, whereas the ionic "spongy" PEDOT:PSS hydrogel slowly squeezes out the $C_{\mathrm{dl}}$ ions.

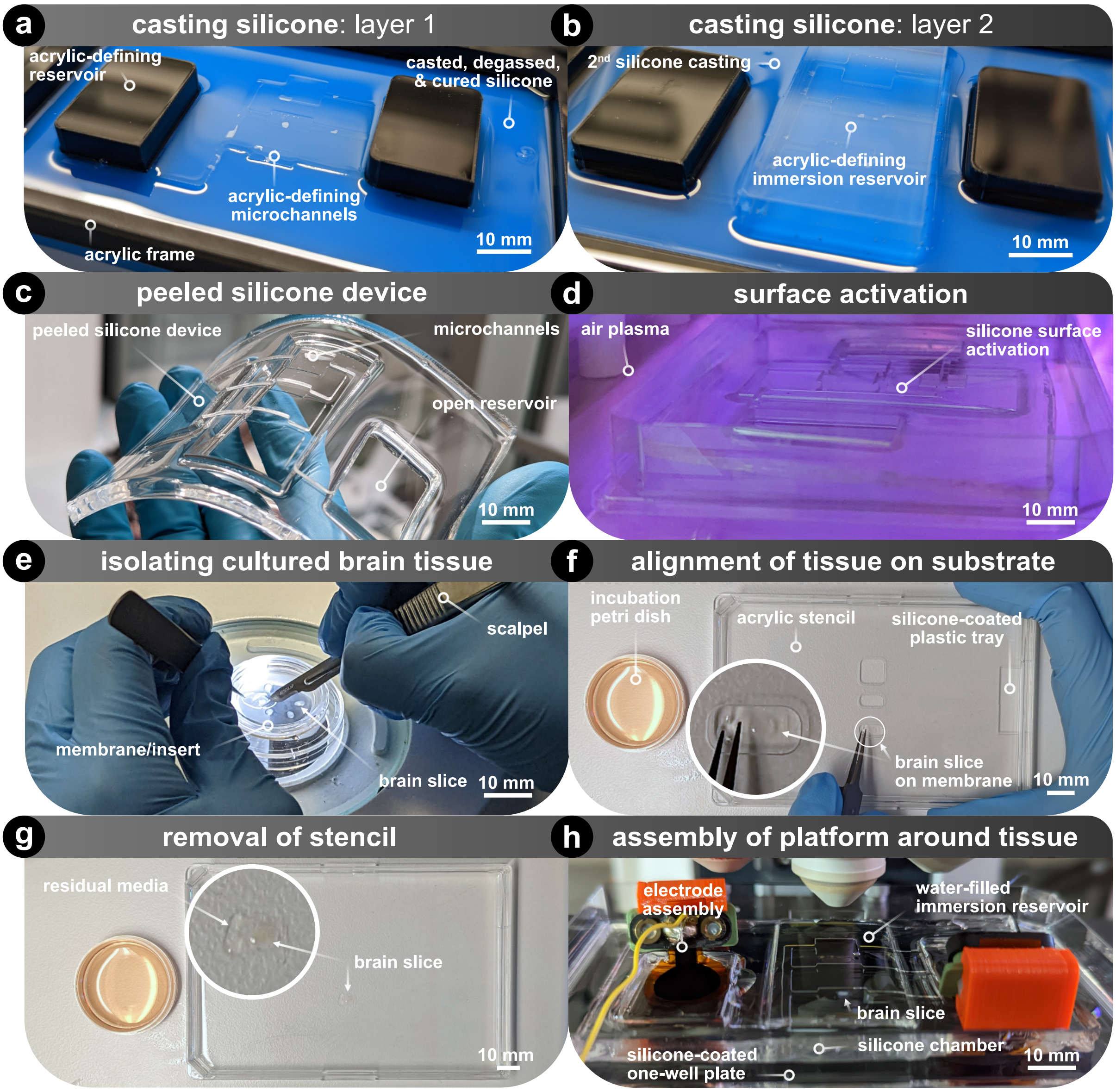


**Fig. S4: Detailed process of how microfluidic chambers are fabricated and assembled**. (a) Acrylic mold is used for casting the first degassed PDMS (silicone) layer in order to define the microchannel, then is cured in the oven at 70 °C for 1 h. (b) More degassed PDMS is added and an acrylic block is added over the cured microchannels in order to create an immersion objective reservoir. Same curing as above. (c) Cured PDMS device is peeled out of the mold. (d) Air plasma treatment (30 W, 30 s, 10 sccm) of PDMS to be temporarily bonded. (e) Organotypic brain slice on membrane insert are cut using a scalpel. (f) Wet brain slice and membrane are transferred to a plasma-treated and silicone-coated one-well polystyrene plate with an acrylic stencil. (g) Acrylic stencil is removed (h) Molded PDMS chamber is hand-aligned and adhered onto the plate. Then, warm media is injected into the reservoir to allow for displacement of air in the microchannels with media. Finally, electrodes are added, water added to the immersion reservoir, and placed on the heated confocal microscope stand.

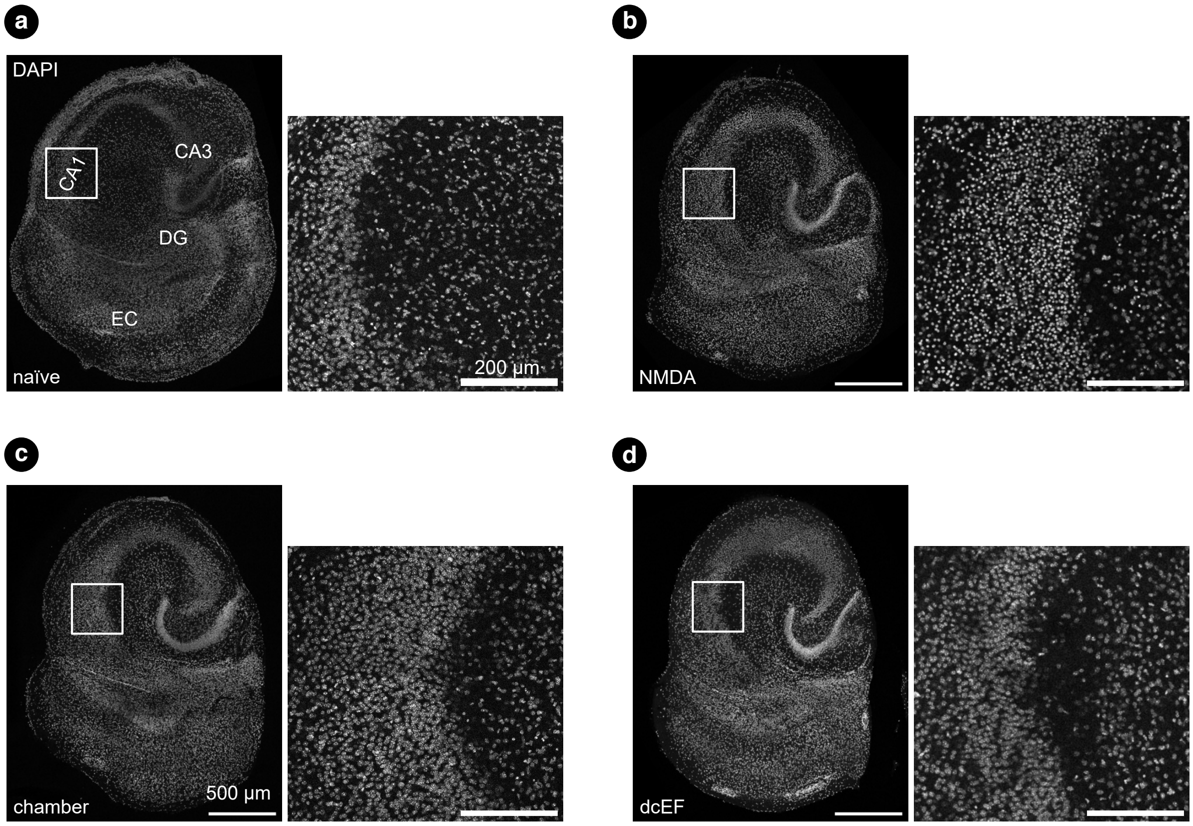


**Fig. S5: DAPI signal of the cultures used in the SYTOX-green experiment**. Experimental design for four groups: naïve controls, positive controls where the cultures were treated with 50 $\mu$M NMDA for four hours, the chamber-immersed group where cultures were immersed inside the chamber for 20 min without switching on DC stimulation, and the dcEF-stimulated group where DC stimulation was applied for 10 min thus cultures in three channels were stimulated correspondingly with three weak EF intensities. The DAPI signal in CA1 cells showed a similar pattern in (a),(c), and (d), but presented a round shape and condensed manner in the positive control group (b), suggesting cell death after NMDA treatment.

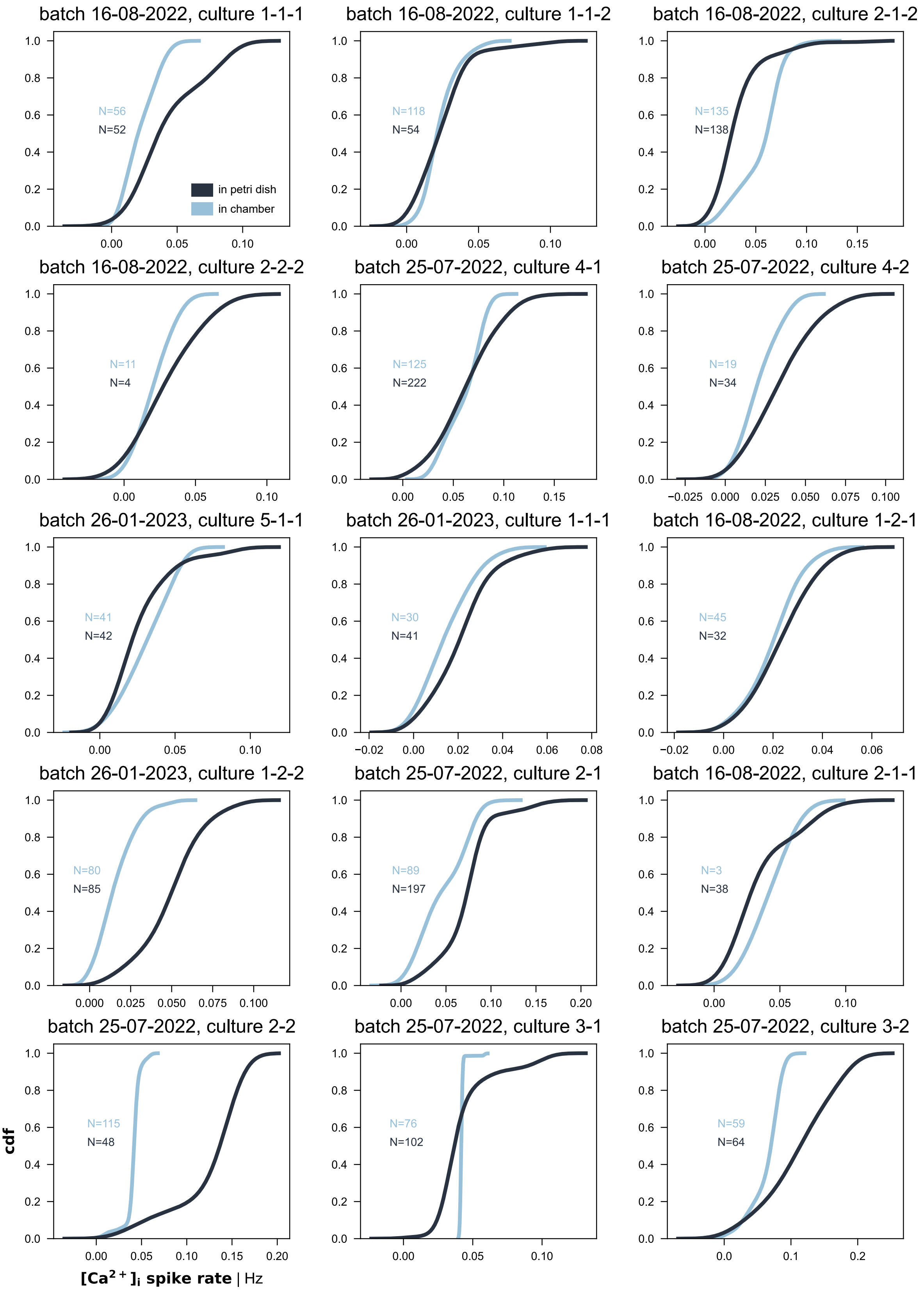


**Fig. S6: Cumulative distribution of calcium spike rate for individual cultures used in Fig 5c.** Within each culture, the calcium spike frequency of individually identified neurons was not all identical but presented a narrow or wide distribution ranging from zero spikes to a few spikes. The distribution varied from culture to culture.

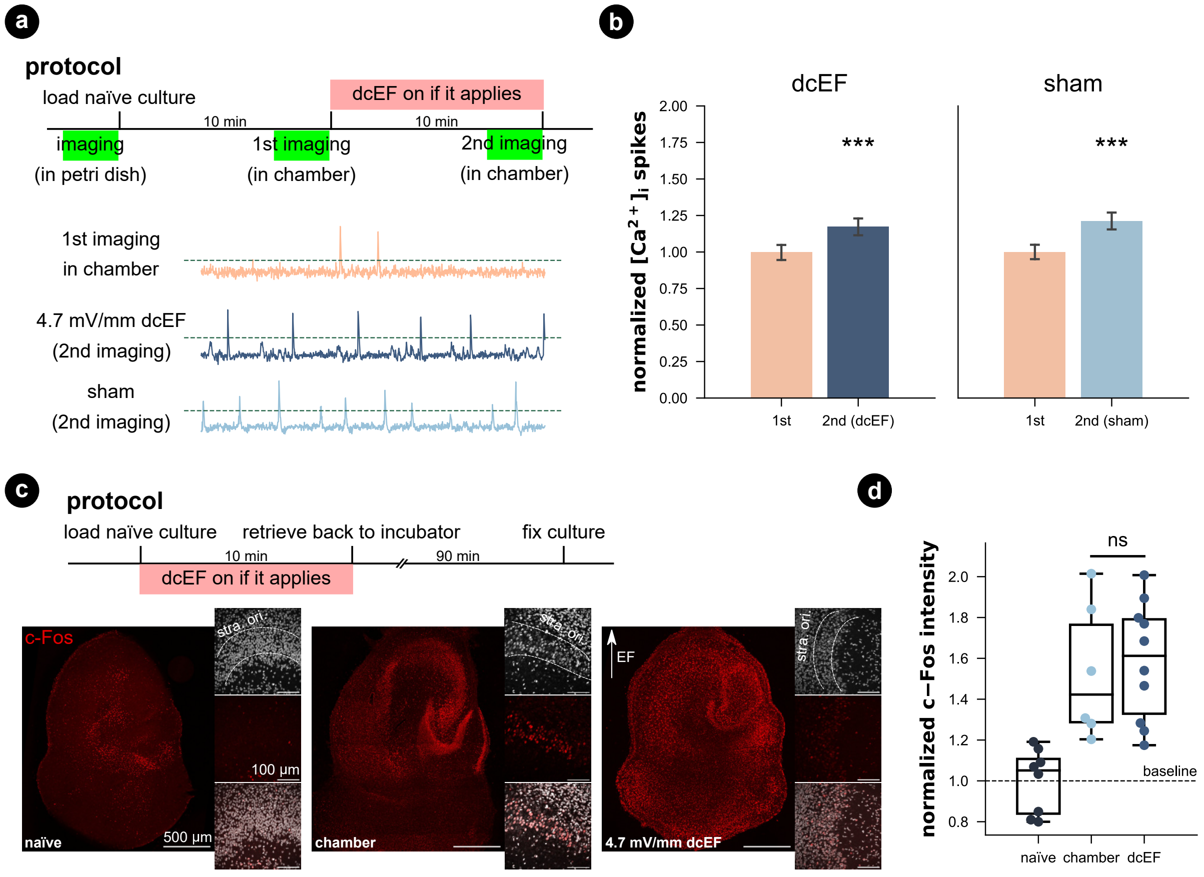


**Fig. S7: Weak dcEF stimulation did not change calcium dynamics or c-Fos expression.** (a) Protocol used for this calcium imaging experiment and sample traces from the corresponding groups. The results comparing the calcium spikes of the same culture in petri dish and in chamber (first two imaging sessions) were summarized in Fig. 5a-c, so it is not displayed here. Then the cultures after the first imaging session were split into two groups in the second imaging session, subject to 4.7 mV $\text{mm}^{-1}$ dcEF stimulation or not (sham). (b) Compared to their corresponding calcium activity levels at the first imaging session inside the chamber, both groups presented significant increases ($p < 0.001$, unpaired student's T-test and justified by LLM where 99.99% CI doesn't cross zero), suggesting a main effect of timing and confirmed with ANOVA and LLM tests. $N = 7$ cultures were used for the sham group (505 neurons identified at baseline and 465 neurons identified for during sham treatment), and $N = 8$ cultures were used for weak dcEF stimulation (497 neurons identified at baseline and 443 neurons identified for during stimulation). Each culture was imaged twice at baseline and under experimentation. The calcium spike rates were normalized by their corresponding average rate at baseline. (c) Protocols used for c-Fos experiment and representative images. (d) Results showed that the handling and immersion inside the chamber (with or without dcEF) increased the c-Fos expression when compared to naïve cultures ($p = 0.0075$ and $p < 0.001$, respectively, Kruskal Wallis test). No difference was observed between chamber-controls and the dcEF-stimulated cultures ($p = 0.87$, Kruskal Wallis test). $N = 8$ for naïve cultures, $N = 6$ for chamber immersed cultures, $N = 10$ for dcEF stimulated cultures.

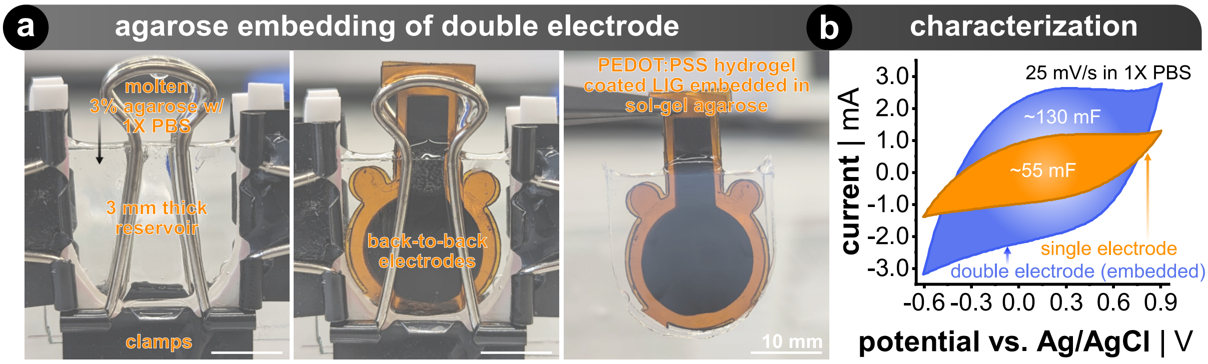


**Fig. S8: Fabrication and CV characterization of a double electrode with salt bridge for pulsed DC current in main text Fig. 5**. (a) A mold was made by sandwiching a laser-cut 3 mm-thick U-shaped acrylic piece between two pieces of white silicone gaskets and glass. This assembly was compressed by clips. The mold was filled with molten 3 % w/v agarose that was dissolved in 1$X$ PBS. The back-to-back electrodes, which each single electrode is the same electrode material and size used throughout the paper, are placed in the molten agarose. After a few minutes the agarose cools down and forms an agarose gel salt bridge that are filled with ions (*e.g.*, $Na^+$ = 157 mM, $K^+$ = 4.45 mM, $Cl^+$ = 142 mM, $HPO_4^{2-}$ = 7.3 mM, and $H_2PO_4^-$ = 4.6 mM). (b) Shows the CV at 25 mV s−1 in 1$x$ PBS for single and double electrodes and their corresponding capacitance.

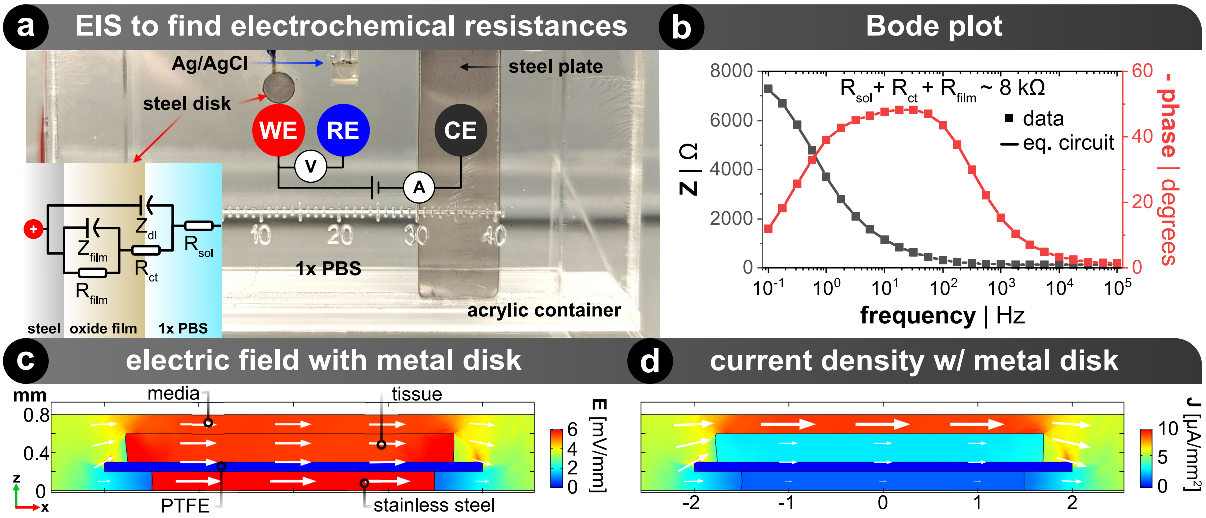


**Fig. S9: Simulation of the EF and current density distribution within the s1 microchannel when metal disk is present.** (a) Three-electrode setup to perform electrochemical impedance spectroscopy (EIS). The same stainless steel disk used in all experiments was used here. A wire was soldered to the back-side and encapsulated with epoxy so that only the front-side of the disk is in contact with the 1× PBS solution. WE is the working electrode, which is the stainless steel disk. RE is the reference electrode, which is a standard Ag/AgCl reference (BASI MF-2052, USA). CE is a large stainless steel sheet. All of which is an acrylic viewing container, same as in Fig. S3. Etched scale bar is in millimeters. The inset diagram shows the equivalent circuit of stainless steel in an electrolyte as described by Rios *et al.*[8] $R_{sol}$, $R_{ct}$, and $R_{film}$ signify the electrolyte solution resistance, the charge transfer resistance, and stainless steel superficial oxide film resistance, respectively. $Z_{dl}$ and $Z_{film}$ represents the constant phase elements of the electrochemical double layer that the electrolyte forms on the metal oxide as well as the coupling between the oxide and bulk steel layers, respectively. (b) EIS Bode plot that shows the stainless steel's electrochemical properties. Of interest is the total resistance for moving ions in solution can transfer its charge into the steel and to transduce the ionic into electronic current within the steel bulk. The sum of the resistances is $\sim$ 8 kΩ, where 73 % comes from $R_{film}$ and 26 % comes from $R_{ct}$. The electrolyte-metal resistivity of stainless steel is approximated by multiplying the aforementioned electrochemical resistance by the electrode area ($0.126\,\mathrm{cm}^2$) and dividing by the distance between WE and CE (15 mm). The reciprocal of resistivity yields the conductivity of ($\sigma = 0.15\,\mathrm{S\,m^{-1}}$), which is $\sim$ 10 x lower than then electrolyte and $\sim$ 3 x lower than the tissue. (c-d) These input conditions correspond with Fig. 6 in the main text, but now with the added metal disk with the aforementioned conductivity value. The slice depicted here is from "s1" in the device. (c) EF distribution confirms a similar EF (14 % increase) within the tissue as compared to when the metal is not present. (d) Current density distribution shows that most of the current travels through the more ionically conductive electrolyte ($\sigma = 1.5\,\mathrm{S\,m^{-1}}$) and electrolyte-filled tissue ($\sigma = 0.5\,\mathrm{S\,m^{-1}}$).

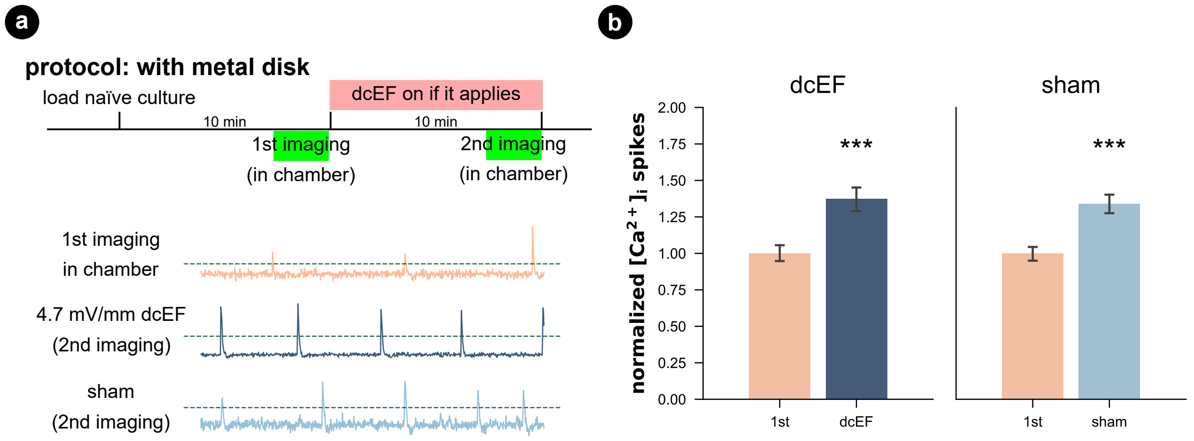


**Fig. S10: Weak dcEF did not change calcium dynamics with the presence of the metal disk.** (a) Protocol used for this calcium imaging experiment was the same as used in Fig. S8 while the cultures were mounted onto a metal disk before being loaded into the chamber and stimulated in the presence of the metal disk. These cultures were not imaged in Petri dish beforehand. After being imaged for the first time, cultures were randomly selected to receive a 4.7 mV mm$^{-1}$ dcEF stimulation or not (sham). The lower panel shows the sample traces from the corresponding groups. (b) Same as the calcium imaging experiments without metal disk, compared to their corresponding calcium activity levels at the first imaging session inside the chamber, both groups presented significant increases ($p < 0.001$, student's T-test and justified by LLM where 99.99% CI doesn't cross zero), suggesting a main effect of timing and confirmed with ANOVA and LLM tests. $N = 6$ cultures were used for the sham group (270 neurons identified at baseline and 305 neurons identified for during sham treatment), and $N = 6$ cultures were used for weak dcEF stimulation on top of the metal disk (442 neurons identified at baseline and 394 neurons identified for during stimulation). Each culture was imaged twice at baseline and under experimentation.

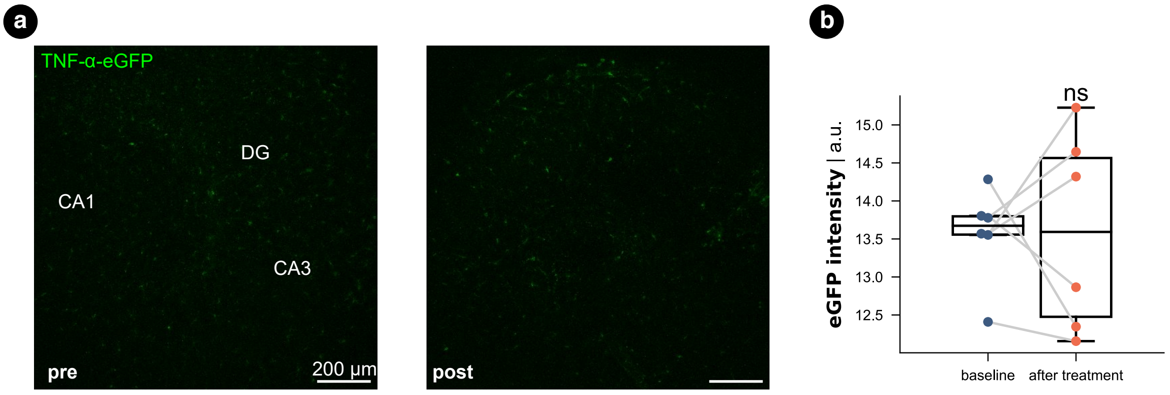


**Fig. S11: Immersion inside the silicone microfluidic chamber with a metal disk did not cause inflammatory-like status in tissue cultures.** (a) Representative images of TNF$\alpha$-eGFP culture before and after 20 min immersion inside the chamber. No dcEF was applied. (b) Quantified eGFP signal intensity of the whole culture before and after immersion. Each dot represents a culture, and the box plots summarize each condition's mean, quartiles, and distribution. $N = 6$ for this experiment. Wilcoxon test was used for statistical analysis. $p > 0.99$.

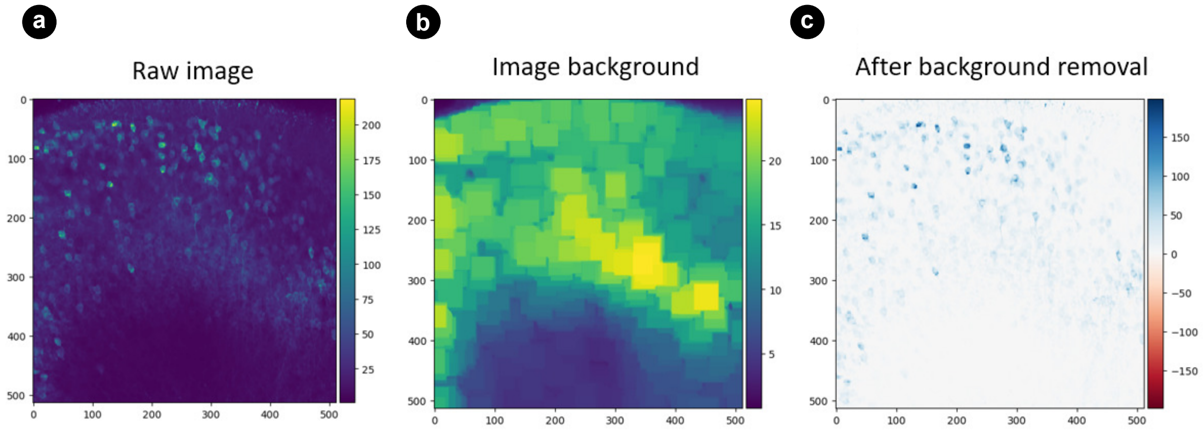


**Fig. S12: Process of detecting neurons for extracting calcium signal time series.**
